# Shank3 mutation disrupts the molecular signature of sleepiness across development

**DOI:** 10.64898/2026.08.21.746326

**Authors:** Elliot Wald, Elizabeth Medina, Caitlin Ottaway, Christine M. Muheim, Kaitlyn Ford, Taylor Wintler Patterson, Kristan Singletary, Ashley Ingiosi, Lucia Peixoto

**Affiliations:** Department of Translational Medicine and Physiology, Sleep and Performance Research Center, Elson S. Floyd College of Medicine. Washington State University, Spokane, WA, United States; Integrative Physiology and Neuroscience, Washington State University, Pullman, WA, United States; Center for Developmental Biology and Regenerative Medicine, Seattle Children’s Research Institute, Seattle, WA, United States; Seattle Children’s Hospital, Seattle, WA, United States; Department of Neuroscience, Ohio State University, Columbus, OH, United States

**Keywords:** Transcriptomics, Autism Spectrum Disorder, Shank3, neurodevelopment, sleep deprivation

## Abstract

Background

Sleep problems are common in autism, emerge early in life and reduce quality of life, yet the mechanistic link between autism and poor sleep remains unclear. Human and rodent data indicate that difficulty falling asleep is a core feature of autistic insomnia, pointing to impaired responses to sleepiness as the underlying cause. We previously showed that adult mice carrying a mutation in the high-confidence autism gene *Shank3* (Shank3^ΔC^) recapitulate this insomnia phenotype and struggle to respond to sleepiness after acute sleep deprivation. Here, we used Shank3^ΔC^ mice to examine the molecular basis of sleepiness and how this autism-associated mutation alters it to inform understanding of sleep problems in autistic individuals.

**Methods:** This study used RNA-sequencing and bioinformatics to identify molecular targets underlying the effect of the Shank3^ΔC^ mutation on the molecular basis of sleepiness across development in male mice. We first compared cortical genome-wide gene expression following acute sleep deprivation and recovery sleep in adult wild-type (WT) and mutant mice. We then used polysomnography and RNA-sequencing to assess the response to increased sleepiness in WT and mutant mice at postnatal days 24 and 30.

**Results:** The neurotypical response to acute sleep deprivation shifted from upregulating neuronal growth and development pathways at P24/P30 to upregulating DNA damage repair and neuronal activity-dependent transcription in adulthood. The Shank3^ΔC^ mutation largely blocked recruitment of these pathways at P24 and in adulthood while paradoxically increasing the magnitude of the mutant response at P30. In addition, mutants consistently upregulated oxidative stress pathways linked to neurodegeneration and protein synthesis regardless of age, whereas WT animals downregulated these functions.

**Limitations:** This study examined gene expression only in male mice, used a single autism rodent model, and averaged signals across mixed cortical cell types. Future work should include females, additional autism models, and single-cell approaches in additional brain regions to further characterize the cellular effects of sleep deprivation and autism-associated mutations.

**Conclusions:** The Shank3^ΔC^ mutation impairs the molecular accumulation of and response to sleepiness, both by elevating oxidative stress responses and by blocking the age-typical upregulation of pathways that differ between juveniles and adults.

## Background

Sleep problems are prevalent in Autism Spectrum Disorder (ASD) and heavily impact quality of life for individuals and caregivers (1). Problems falling asleep can be detected in infancy before ASD diagnosis and are associated with altered patterns of brain development, and difficulty falling asleep remains a core feature of autistic insomnia in school-age children (2,3). We have previously shown that adult mice with a mutation in the high-confidence ASD gene *Shank3* (Shank3^ΔC^) recapitulate the ASD clinical sleep phenotype (4,5). Adult Shank3^ΔC^ mice are unable to increase sleep amounts and have long sleep onset latencies in response to acute sleep deprivation (SD) (4,5). In addition, our studies on young Shank3^ΔC^ mice show that the Shank3^ΔC^ mutation prevents the development of the ability to fall asleep faster in response to increased sleep pressure between postnatal days 24 (P24) and 30 (P30), resulting in animals that take a long time to fall asleep after SD regardless of age (6). These studies suggest that the Shank3^ΔC^ mutation interferes with the development of sleep homeostasis: the process of balancing time spent awake through a compensatory increase in sleep time and quality. However, the molecular mechanisms underlying sleep homeostasis and how the Shank3^ΔC^ mutation may affect them remain poorly understood.

Multiple studies have used transcriptomics following acute SD to define molecular processes that underlie sleep homeostasis at the cellular level. Acute SD in adult mice leads to an upregulation of immediate early genes (e.g. *Homer1a*, *Fos*, *Arc*, *Bdnf*), pathways associated with neuronal activity (e.g. MAPK and Ras signaling) and circadian rhythms (7–9), recruitment of the unfolded protein response and changes in macromolecule biosynthesis (10,11) as well as the downregulation of pathways involved in protein synthesis associated with deficits in learning and memory (12–14). Additionally, acute SD has been shown to affect these processes in predominantly glutamatergic neurons (15). However, the pathways that track sleep need, rather than being a non-specific response to SD (such as stress), are less understood. By matching transcriptomic profiles to sleep pressure dynamics (different amounts of SD and subsequent recovery sleep) we identified redox metabolism, chromatin regulation and DNA damage/repair as molecular mechanisms linked to sleep homeostasis in adult mice (16). In addition, we showed that the transcriptional response to SD changes dramatically during postnatal development in mice, mainly affecting developmental signaling (such as Wnt signaling) in juveniles and biomolecular/metabolic pathways in adults (17). Whether mutations associated with ASD impair the recruitment of the above-mentioned pathways in response to increased sleep need and whether that effect changes as sleep homeostasis regulation develops postnatally remains unexplored.

The objective of this study was to use transcriptomics to define how the Shank3^ΔC^ mutation affects molecular pathways in response to sleep need across development in the male mouse frontal cortex, with the goal of identifying molecular mechanisms linking Shank3 and sleep homeostasis. We previously showed that SD increases gene expression differences between wild-type (WT) and Shank3^ΔC^ with a particular effect on circadian transcription factors in the cortex of adult male mice (4). However, how the mutation affects dynamics of gene expression in response to increased sleep need (acute SD and subsequent recovery sleep), as well as how this response may change in juveniles versus adults, remains unknown. We found that the molecular response to acute SD changes across development in WT animals but remains similar in Shank3^ΔC^ animals, with mutants showing increased oxidative stress following SD while simultaneously being unable to mount a typical WT response. In WT mice, the response to SD shifts from an upregulation of neuronal growth and development pathways at P24 and P30 to an upregulation of intracellular signaling, DNA damage/repair and long-term potentiation in adulthood. This shift does not occur in Shank3^ΔC^ animals. Instead, mutants upregulate oxidative stress pathways linked to neurodegeneration and protein synthesis/processing after SD regardless of age, molecular mechanisms that WT animals downregulate. These findings suggest that the Shank3^ΔC^ mutation may impair sleep homeostasis at the molecular level in two ways: by preventing the recruitment of developmentally appropriate pathways in response to sleep need and by causing an aberrant over-activation of oxidative stress pathways regardless of age.

## Methods

### Experimental design and animals

Male Shank3^ΔC^ and WT C57BL/6J littermate mice at three ages, postnatal day 24 (P24), P30 and P90 (adult, 10-12 weeks), were used. Briefly, animals were individually housed in standard cages at 24 ± 1°C on a 12:12 hour light/dark cycle with food and water *ad libitum* for 6-7 days before tissue collection. Adult mice were assigned to sleep deprivation (SD), SD and recovery sleep (SD+RS), and allowed-to-sleep home cage (HC) groups, n = 5 per genotype and condition. P24 and P30 mice were assigned to SD and HC groups, n = 5 per genotype and condition. Beginning at light onset (zeitgeber time (ZT) 0), SD mice were kept awake for either 3 hours (P24, P30) or 5 hours (adult) via gentle handling to maximize sleep drive as outlined in our previous work (17). SD+RS animals (adult only) were given a 2-hour recovery sleep opportunity after 5 hours of SD. HC mice served as time-matched controls and were left undisturbed in their home cage for either 3, 5, or 7 hours beginning at light onset. All experimental procedures were approved by the Institutional Animal Care and Use Committee of Washington State University and conducted in accordance with National Research Council guidelines and regulations for experiments with live animals.

### Tissue collection, RNA isolation, library preparation and sequencing

Frontal cortex tissue was homogenized in Qiazol buffer (Qiagen, Hilden, Germany) using a TissueLyser (Qiagen) and all RNA was extracted using the Qiagen RNAeasy kit (Qiagen) on the same day. RNA-sequencing library preparation, quantification and sequencing was performed at WSU Spokane Genomics Core. The integrity of total RNA extracts was assessed using a Fragment Analyzer (Advanced Analytical Technologies, Inc, Ankeny, IA) with the High Sensitivity RNA Analysis Kit (Advanced Analytical Technologies, Inc). High quality samples were used for RNA library preparation with the TruSeq Stranded mRNA Library Prep Kit (Illumina, San Diego, CA). Briefly, mRNA was isolated from 2.5 mg of total RNA using poly-T oligo attached magnetic beads and then subjected to fragmentation, followed by cDNA synthesis, dA-tailing, adaptor ligation, and PCR enrichment. RNA library sizes were assessed by a Fragment Analyzer with the High Sensitivity NGS Fragment Analysis Kit (Advanced Analytical Technologies Inc). The concentrations of RNA libraries were measured by a StepOnePlus Real-Time PCR System (ThermoFisher Scientific, San Jose, CA) with the KAPA Library Quantification Kit (Kapa Biosystems, Wilmington, MA). Paired-end sequencing was performed using HiSeq 2500 SBS kit v4 (Illumina) with a read length of 100 bp and an average sequencing depth of 78 million reads per sample. HiSeq Control Software (v. 2.2.68) was used for base calling, and raw bcl files were converted to FASTQ files using bcl2fastq (v. 2.17.1.14). Adapters were trimmed from the FASTQ files during the conversion.

### Transcript quantification

Raw sequencing reads were processed with Salmon (v. 1.10.0) using default parameters (18). To improve the accuracy of quantification estimates from Salmon, an index was built that incorporated a set of genome targets as decoys using the reference genome and transcriptome annotation files from GENCODE (release M37, primary assembly GRCm39, Ensembl release 114) as previously described (15). Quantification was completed with ‘salmon quant’ using the following parameters ‘--threads 6’, ‘-l À, and ‘--numBootstraps 30’.

R/Bioconductor (v. 3.20.0) packages were used during the downstream analysis pipeline. Tximeta (v. 1.24.0) was used to summarize to the gene level (19).

### Differential expression analysis

Gene counts were first filtered to avoid zero inflation using the Fishpond package (v 2.12.0). Genes with less than ten reads across five samples were filtered out of the data set. Statistical analysis was performed for the juvenile (P24/P30) and adult animals independently. Upper-quartile normalization using the EDASeq package (v. 2.40.0) was performed first to correct for differences in library size (20) and RUV was subsequently used to remove additional unwanted variation using replicate samples (v. 1.40.0) (21). Parameter k = 15 was used for both the P24/P30 data and the adult data. Differential gene expression was estimated using the nonparametric expression analysis method from Fishpond Swish (v. 2.12.0) (22). The FDR cutoff was < 0.05. Within each age, SD animals were compared to HC animals of the same genotype. The following differential expression comparisons were done relative to HC controls independently for WT and Shank3^ΔC^ mice: HCP24 vs. SDP24, HCP30 vs. SDP30, HCP90-5 vs. SDP90 and HCP90-7 vs. SD+RSP90 for a total of 8 comparisons. For each comparison, differentially expressed genes were visualized by plotting the log2 fold changes of all expressed genes in WT versus Shank3^ΔC^ mice and color coding based on the FDR cutoff and group specificity. Positive control genes were assembled from a previous study (16) based on adult male wild-type mice and were used to evaluate the reproducibility of differential gene expression for adult wild-type samples only. Control gene probe IDs were mapped to Ensembl IDs then annotated using Ensembl release 114 (23) using the package biomaRt (v. 2.62.1).

### Prioritization and functional enrichment analysis

To further focus our analysis, lists of total differentially expressed genes were intersected and visualized using the UpSetR (v. 1.4.0) and ComplexUpset (v. 1.3.3) packages from R/Bioconductor to define all possible intersections between timepoints and genotypes for upregulated and downregulated lists. Intersections that represented unique genes (genes only associated with one genotype and timepoint) were selected for further analysis.

Functional annotation of the chosen gene lists was performed using the Database for Annotation, Visualization, and Integrated Discovery v2021 (DAVID) with DAVID Knowledgebase v2025_1 (24). The UniProt terms biological process (BP) and molecular function (MF) along with the Kyoto Encyclopedia of Genes and Genomes (KEGG) pathways were selected as the annotation categories. For each functional term, enrichment was defined relative to all expressed genes in the frontal cortex (21,852 after filtering) using EASE score <0.05. All enriched terms were clustered based on overlap of genes using final group memberships of three and a similarity threshold of 0.20 (at least 20% overlap in genes). Both clustered and unclustered enriched terms were visualized using ggplot2 (v. 3.5.2). Hub genes were defined by identifying genes that appeared in all or almost all the individual terms for a functional annotation cluster.

### EEG data analysis

The sleep homeostatic response primarily manifests itself as an increase in sleep time and continuity early in life (25–28). We had previously not detected changes in sleep pressure from WT and Shank3^ΔC^ mice following SD at P24 and P30 (6), so we reanalyzed our lab’s publicly available sleep recording data to determine if the mutation led to differences in sleep time after acute SD. Data analysis, visualization, and statistics were performed using MATLAB (MathWorks). Time in state (Wake, non-rapid eye movement sleep- NREM- and rapid eye movement sleep-REM) was measured as a percentage of total recording time (TRT) during the light (ZT 0-12) and dark periods (ZT 13-24) and displayed as the mean ± standard error of the mean (SEM) for each state. Analysis of time in state was performed independently for the light period and dark periods because of the known differential effect of light on sleep parameters (29). Repeated measures ANOVA using time (light period ZT 3-12 or dark period ZT 13-24) as the repeated measure and treatment day (baseline or SD) as the between subjects’ variable was used to determine significant differences in time in state between baseline and SD within genotype.

### Data availability

Sequencing data are available through the Gene Expression Omnibus Database (GEO) under accession number GSE306621. Data from wild-type animals used in this study were obtained from GSE211301 (SD and HC, P24, P30) and GSE113754 (SD and HC, adult). Raw EEG data are available at https://sleepdata.org/datasets/medina-2022. All code used for analysis and visualization is available on Github: https://github.com/PeixotoLab/RNAseq_sleep_developmentShank3

## Results

The goal of this study was to use genome-wide gene expression to uncover molecular processes and pathways that may mediate the inability of Shank3^ΔC^ mice to mount a typical sleep homeostatic response across postnatal development. To do so, we used RNA-seq in the mouse frontal cortex following acute SD and subsequent recovery sleep (SD+RS) across varying ages in male Shank3^ΔC^ animals and their WT littermates to better understand how an ASD-associated mutation may disrupt sleep homeostasis at the molecular level across postnatal development and into adulthood.

### The Shank3^ΔC^ mutation alters transcriptional dynamics following sleep deprivation and subsequent recovery sleep in adult animals

We first examined how gene expression dynamics after sleep deprivation (SD) and subsequent recovery (SD+RS) were affected by the Shank3^ΔC^ mutation in adulthood to identify genes and pathways that were specifically responsive to changes in sleep need. Our results show that the Shank3^ΔC^ mutation impairs genome-wide differential gene expression dynamics following both SD and SD+RS (Figure 1). Figure 1A depicts the log2 fold change (logFC) of all expressed genes in WT and Shank3^ΔC^ mice after SD (as compared to respective undisturbed HC control animals). There are 5719 differentially expressed genes (DEGs, shown in grey) that are shared in both WT and Shank3^ΔC^ animals following SD which include some immediate early genes (IEGs: *Fos*, *Arc, Egr2*) as well as genes involved in the unfolded protein response (*Hspa1b*, *Hspa5*). Interestingly, WT animals have substantially more genotype-specific DEGs (3407, shown in black) with larger logFC than Shank3^ΔC^ animals (1724, shown in red). SD-specific DEGs in WT animals include genes with overlapping roles in gene expression regulation and synaptic plasticity (*Crebbp, Prkcb*), DNA damage/repair (*Ruvbl2*), protein synthesis and regulation (*Eif4e/2, Rps26, Rpl19, Psmd3, Psmc3*), insulin signaling (*Pik3r3, Eif4e/2)* and the IEG *Junb*. Figure 1B depicts the logFC of all expressed genes in WT versus Shank3^ΔC^ after SD+RS. In this case, Shank3^ΔC^ animals have more DEGs after RS than WT (2612 vs 2203). In WT animals, the difference in DEGs between SD (5719 shared + 3407 unique) and SD+RS (4280 shared + 2203 unique) shows that RS leads to a large reduction (>2500) in the number of gene expression changes as we have previously shown (16). Unlike WT animals, Shank3^ΔC^ mutants only display an ∼500 gene difference in the number of DEGs after SD (5719 shared + 1724 unique) and SD+RS (4280 shared + 2612 unique). Overall, our data demonstrates that the Shank3^ΔC^ mutation dramatically inhibits the number of DEGs that respond to SD and reduces the impact of RS on the recovery of gene expression changes after SD. Additional File 1 contains the lists of total DEGs for adult animals. Additional File 2 shows data processing and normalization quality control plots (Additional File 2 A, B) as well as minus-average (MA) plots depicting the log average vs log ratio of gene expression for each of the pairwise comparisons in adult animals (Additional File 2 C-F).

**Figure 1.**
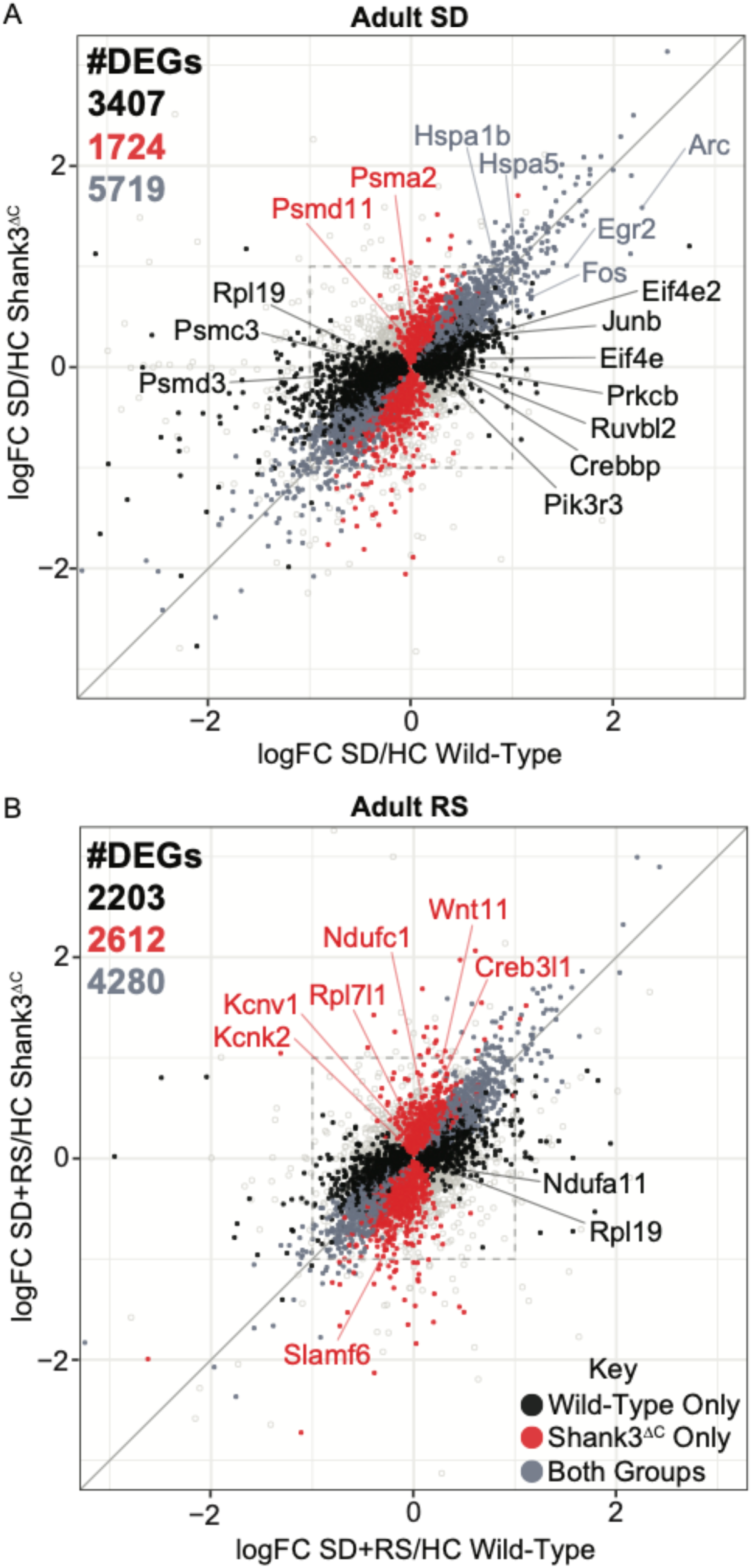
The Shank3^ΔC^ mutation impairs gene expression dynamics following SD and subsequent RS in adults. Correlation plot between the logFC of DEGs in adult WT (x-axis) and Shank3^ΔC^ (y-axis) animals following **A**) SD/HC and **B**) SD+RS/HC. DEGs found in both WT and Shank3^ΔC^ are shown in grey, DEGs unique to WT are in black and DEGs unique to Shank3^ΔC^ are in red. Plot windows are adjusted to show genes with a logFC ≤ |3|. The box represents a logFC cutoff of ± 1. Select genes of interest are highlighted.

To identify genes (and subsequently pathways) that may mediate the effect of the Shank3^ΔC^ mutation on the molecular homeostatic response to SD, we first defined DEGs that were unique to SD or SD+RS in both genotypes by intersecting upregulated and downregulated DEGs for each group (Figure 2). This intersection allows for the identification of genes that are differentially expressed after SD but recover during RS which indicates that they respond specifically to the accumulation and discharge of homeostatic sleep pressure (16). Additionally, it allows us to identify genes that are unique to RS and therefore could have functions that help make RS restorative (16). Figure 2A displays upregulated genes (shown in green) while Figure 2B displays downregulated genes (shown in purple). WT animals upregulate 820 unique genes, while the Shank3^ΔC^ mutants upregulate 454 unique genes after SD. After SD+RS, WT animals upregulate 591 unique genes while mutants upregulate 544. A similar pattern is seen in the downregulated genes, with 1443 genes unique to WT after SD and only 500 unique to mutants. After RS, 605 downregulated genes are unique to WT, while 567 are unique to mutants. This pattern mirrors that observed in Figure 1, where the Shank3^ΔC^ mutation inhibits the molecular homeostatic response to SD. A complete list of DEGs for adults is provided in Additional File 3.

**Figure 2.**
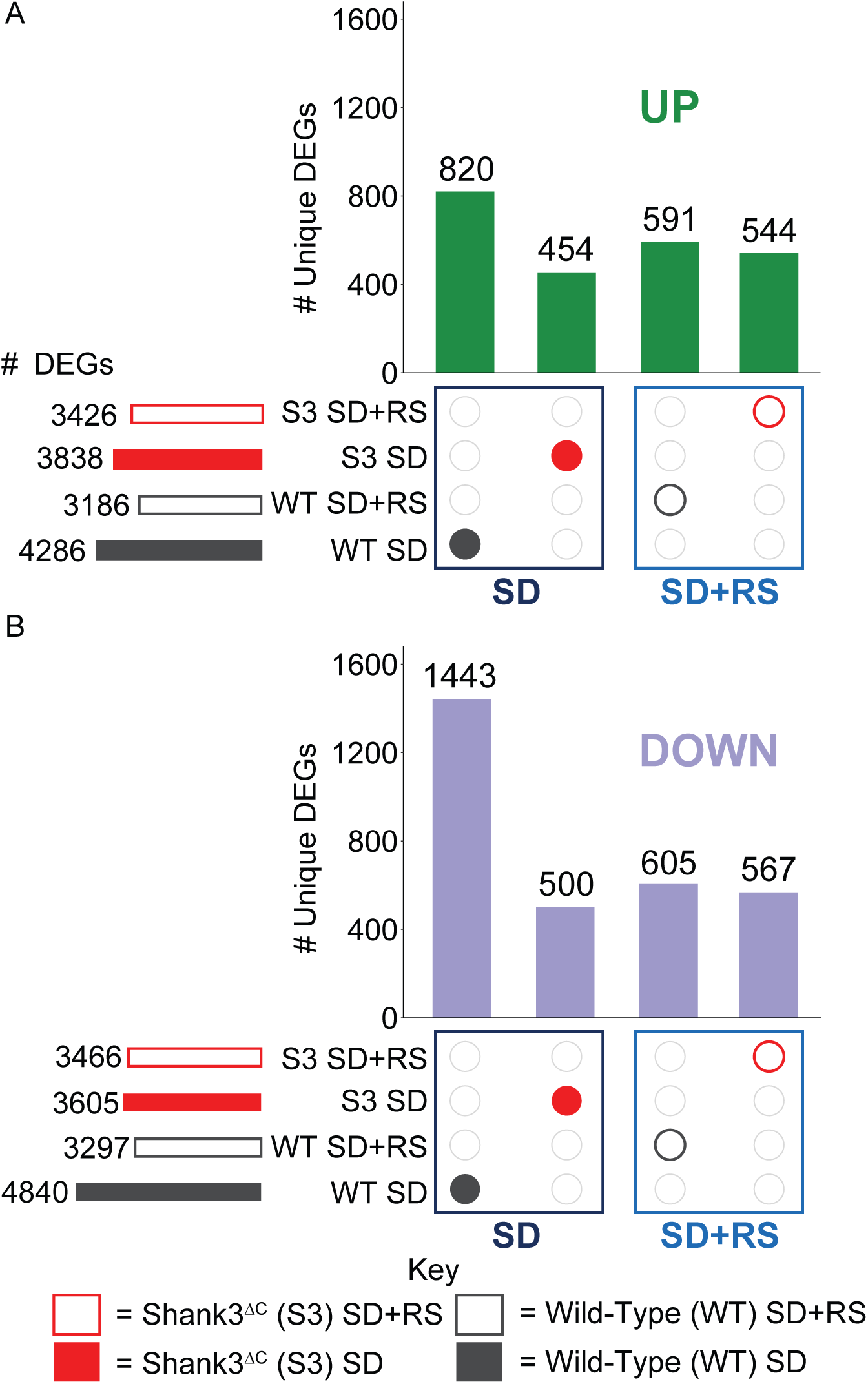
Adult WT mice have increased differential gene expression following SD, RS. UpSet plots of the intersections of the number of DEGs between adult Shank3^ΔC^ (red) and WT (dark grey) animals’ responses to SD (navy blue box) and SD+RS (blue box). Lists of DEGs were intersected for **A**) upregulated DEGs (green) and **B**) downregulated DEGs (purple). The total number of DEGs for each condition are shown in the set size rows on the left of the plot. The vertical bars represent the number of DEGs unique to the subset indicated by the colored dots in the intersection matrix. Filled dots represent the SD condition and outlined dots represent the SD+RS condition. Intersections selected for functional enrichment analysis are shown.

To better understand which pathways, molecular processes and biological functions (herein referred to as “functions”) distinguish the WT vs mutant response to SD and RS, we performed functional enrichment analysis of genotype- and condition-specific DEGs. We first identified genes whose expression changed after SD but were no longer differentially expressed after RS (Figure 3). Figure 3A shows functions enriched in DEGs unique to SD in WT animals. Our results indicate that most upregulated functions form one cluster of intracellular signaling and long-term potentiation pathways (*Prkcb, Rac1, Crebbp, Rap1b)*. Additional functions include DNA damage/repair and transcriptional regulation pathways (*Ruvbl2, Rad51c, Atm, Junb*). Downregulated functions form two clusters: ribonucleo/ribosomal protein pathways (*Rpl19, Rps26, Mrps21)* and pathways related to neurodegenerative diseases such as prion, Parkinson’s, and Huntington’s (*Psmc3, Psmd3*). Additional downregulated functions include metabolic and muscle-related processes (*Pgm2, Gpl1*) as well as immune signaling pathways (*C1gtnf4*). Figure 3B shows functions enriched in DEGs after SD unique to Shank3^ΔC^ mutants. Mutants show one cluster of upregulated genes involved in neurodegenerative disease-related pathways such as prion, Parkinson’s, Huntington’s, Alzheimer’s disease and Amyotrophic lateral sclerosis (*Psma2, Psmd11, Itpr1, Mapk14, Wnt4, Cox7b*). Additional functions include upregulation of transcription regulation and signaling (*Eif3, Ep300*) and downregulation of DNA binding and immune-related signaling. Overall, the Shank3^ΔC^ mutation leads to a sharp reduction in biological functions and pathways that SD recruits and RS recovers. In addition, SD leads to the upregulation of neurodegenerative pathways in mutants while these same functions are downregulated in WT after SD. Additional File 4 contains the annotated results of the DAVID analysis depicted in Figure 3.

**Figure 3.**
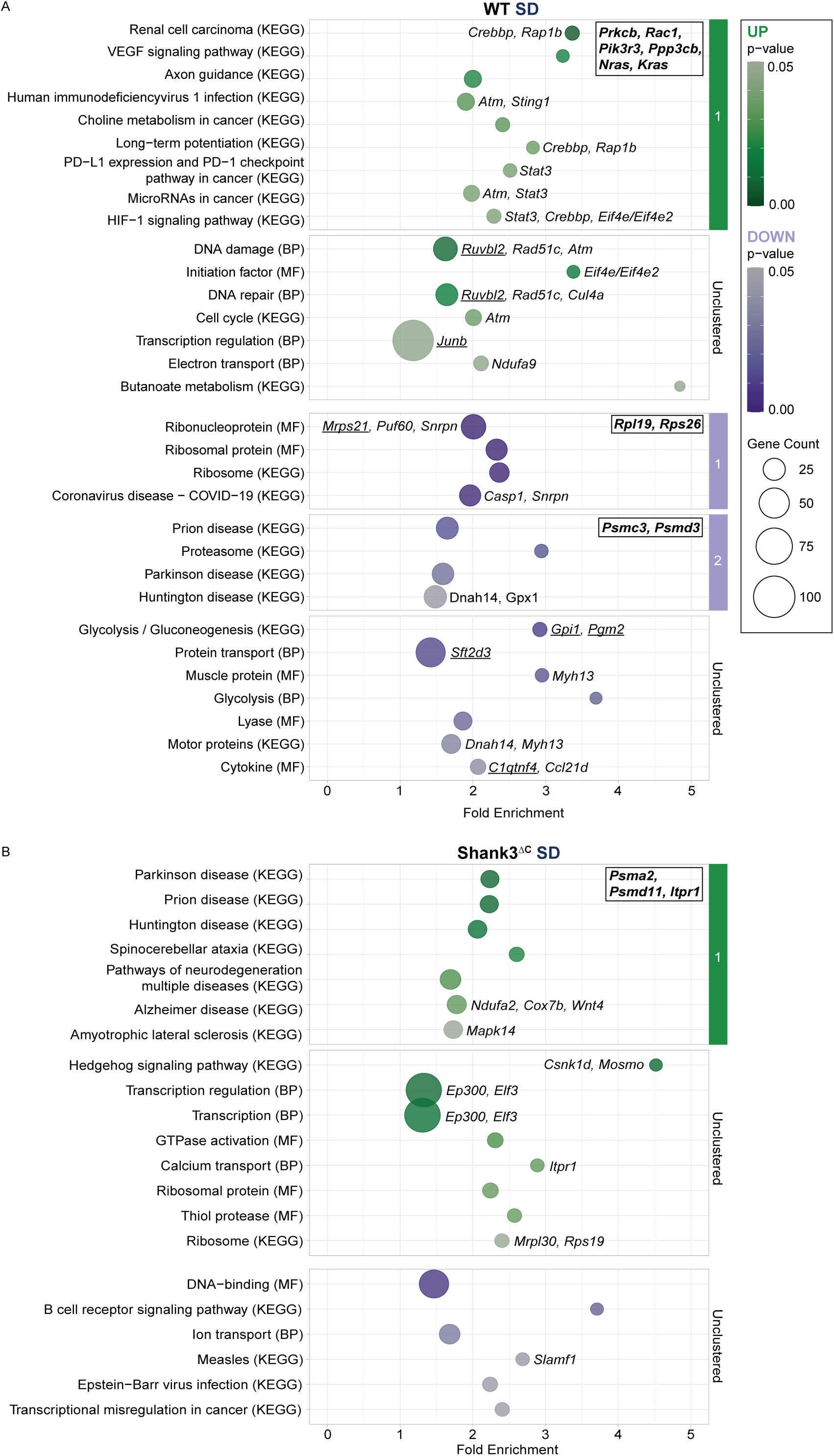
SD upregulates LTP and DNA damage repair in adult WT but not Shank3^ΔC^ animals. Functional enrichment analysis plots for DEGs unique to 5 hours of SD in adult **A**) WT animals and **B**) Shank3^ΔC^ animals. Enriched functional annotation terms (modified Fisher’s exact *P* value < 0.05) from UniProt BP, MF and KEGG are displayed vertically. Circle size indicates the number of genes found within a given term which are plotted based on fold enrichment as displayed on the x-axis. Upregulated terms are in green and downregulated terms are in purple with darker shades representing smaller *P* values. Hub genes are bolded and displayed in boxes in the upper right corner of their corresponding cluster. Enrichment scores for each cluster in **A**: upregulated cluster 1 (1.59), downregulated cluster 1 (3.15), downregulated cluster 2 (1.62). Enrichment scores for each cluster in **B**: upregulated cluster 1 (1.70). Positive control genes are underlined.

Next, we focused on genotype-specific DEGs that were not differentially expressed after SD but induced following RS (Figure 4). Figure 4A shows functions enriched in DEGs after SD+RS unique to WT. Upregulated functions form two clusters, one of ribosome formation pathways (*Rpl19, Rps26, Rps9*) and a second of neurodegenerative disease-related pathways such as Prion, Parkinson’s, Huntington’s, Alzheimer’s disease and Amyotrophic lateral sclerosis (*Ndufa11, Ndufs7, Uqcr11*). These are the same functions that were initially downregulated after SD in WT. Additional upregulated functions include chromatin remodeling and protein synthesis (*Ruvbl1, Macroh2a2, Nsun2*). Downregulated DEGs unique to WT after RS form one cluster consisting of immune and inflammatory signaling and cellular senescence (*Mapk12, Mapk14, Raf1*). Additional downregulated functions include metabolic and cellular regulation (*Galntl6, Pten, Cul4a)*. Figure 4B shows functions enriched in DEGs after SD+RS unique to mutants. Even though mutants have a larger number of genotype-specific DEGs than WT following SD+RS, there is only one upregulated cluster consisting of hormone signaling and endocrine processes (*Sp1, Creb3l1, Creb3l4*). Other upregulated functions include potassium channel activity (*Kcnv1, Kcnk2*) and developmental pathways (*Wnt11, Sox2*). Downregulated functions were primarily associated with adaptive immunity (*Slamf6, Fyn, Myo1g*). These results show that, while the WT animals show a dynamic response to RS with the upregulation of previously downregulated functions, the Shank3^ΔC^ mutants recruit fewer biological pathways during RS. Additional File 5 contains the annotated DAVID output that is represented in Figure 4.

**Figure 4.**
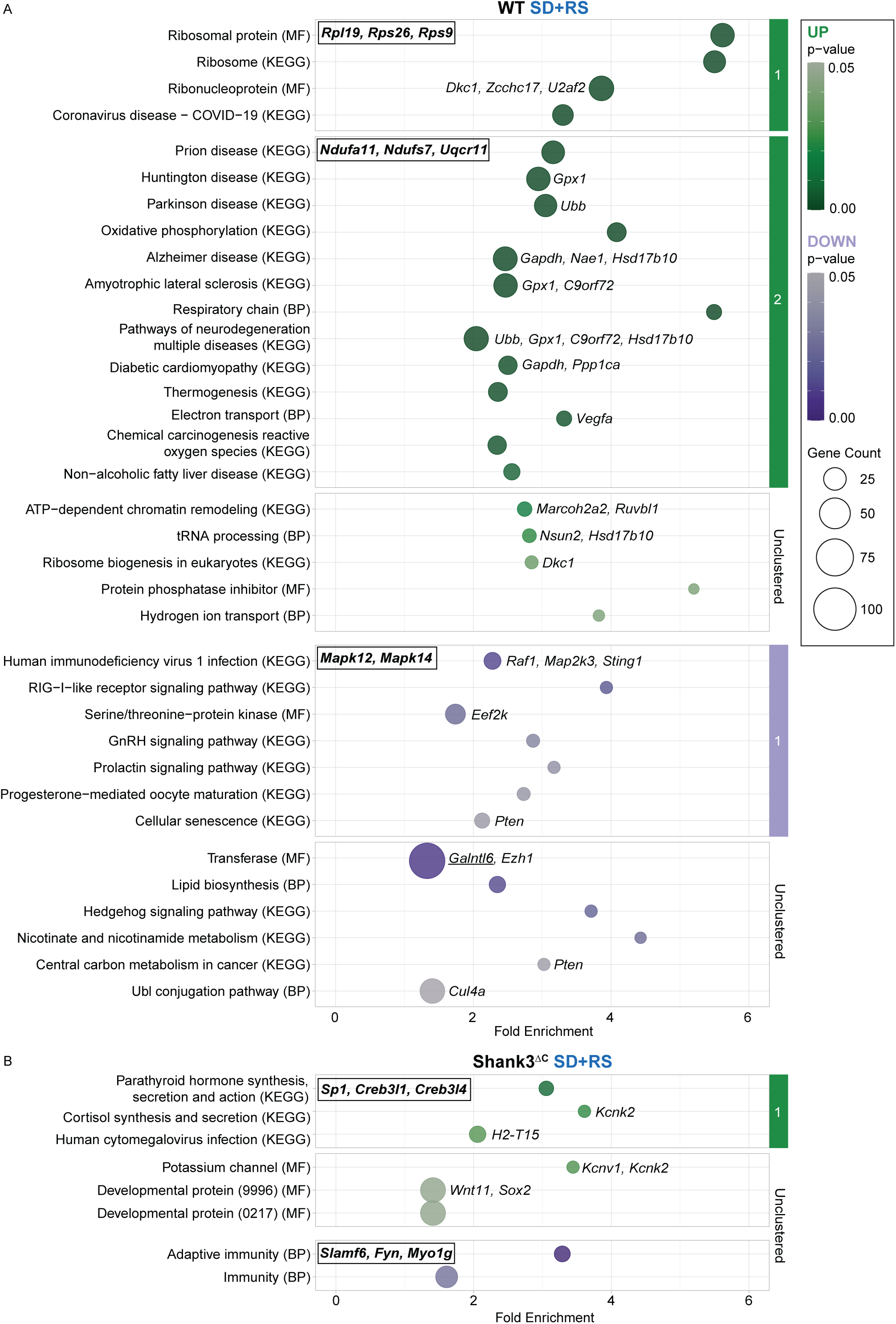
In WT animals, functions unique to SD+RS compensate for the molecular changes caused by SD. Functional enrichment analysis plots for DEGs unique to 2 hours of RS in adult **A**) WT and **B**) Shank3^ΔC^ animals. Enriched functional annotation terms (modified Fisher’s exact *P* value < 0.05) from UniProt BP, MF and KEGG are displayed vertically. Circle size indicates the number of genes found within a given term which are plotted based on fold enrichment as displayed on the x-axis. Upregulated terms are in green and downregulated terms are in purple with darker shades representing smaller *P* values. Hub genes are bolded and displayed in boxes in the upper left corner of their corresponding cluster. Enrichment scores for each cluster in **A**: upregulated cluster 1 (9.44), upregulated cluster 2 (4.13), downregulated cluster 1 (1.56). Enrichment scores for each cluster in **B**: upregulated cluster 1 (1.73). Positive control genes are underlined.

Lastly, we identified the molecular response to SD and SD+RS that was not affected by the Shank3^ΔC^ mutation (Additional File 6). We identified 1459 DEGs unique to SD (784 upregulated and 675 downregulated) and 614 DEGs unique to SD+RS (278 upregulated and 336 downregulated) that were shared between WT and Shank3^ΔC^ mice. The lists of DEGs for common intersections can be found in Additional File 7 and the results of functional enrichment analysis to determine the functions of DEGs not affected by the mutation are shown in Additional File 8. Additional File 8A depicts functions unique to SD but common to mutants and WT. There are two clusters in 784 upregulated genes: transcriptional regulation (*Bhlhe40, Parp1, Per1*) and inflammatory responses/insulin signaling (*Ddit4, Depp1, Nfkbia, Pik3ca*). Additional upregulated functions are circadian rhythms (*Per1, Bhlhe40*) and Ubl conjugation (*Egr2, Kctd21*). The 675 downregulated genes unique to SD in both genotypes are enriched in functions including G-protein coupled receptors (*S1pr5*) and transducers (*P2ry2*). Additional File 8B depicts functions unique to SD+RS but common to mutants and WT. For the 278 upregulated DEGs, there is one upregulated cluster related to ribosome function (*Mrpl26, Mrpl21*) and no downregulated clusters. The full set of functional analysis results is shown in Additional File 9. As these groups of genes and functions are not affected by the mutation, we considered them unlikely to drive the genotype difference in sleep homeostasis we have previously described (4,5) and did not consider them in subsequent analyses.

### The Shank3^ΔC^ mutation alters the transcriptional response to SD during critical periods of development

Early in life, the sleep homeostatic response is known to differ from that in adulthood, and is mainly defined by an increase in sleep amounts (25–28). Although we have shown that male Shank3^ΔC^ mice have abnormal sleep as early as P23, whether the Shank3^ΔC^ mutation leads to an inability to increase sleep amounts in response to SD in young mice remains an open question (6). To address this, we analyzed sleep recordings from our previously published study of WT and mutant mice following SD at P24 and P30. Figure 5 shows that at both P24 and P30, WT animals have a significant increase in non-rapid eye movement (NREM) sleep after SD (WTP24 NREM, light period, p = 0.028; WTP30 NREM, dark period, p = 0.045) while mutants do not. Overall, our data demonstrates that sleep homeostasis deficits are present in juvenile Shank3^ΔC^ mutants as young as P24, while a deficit in latency to sleep onset after SD arises at P30 (6). Time in state for wake and REM can be found in Additional File 10, and full statistical analysis can be found in Additional File 11.

**Figure 5.**
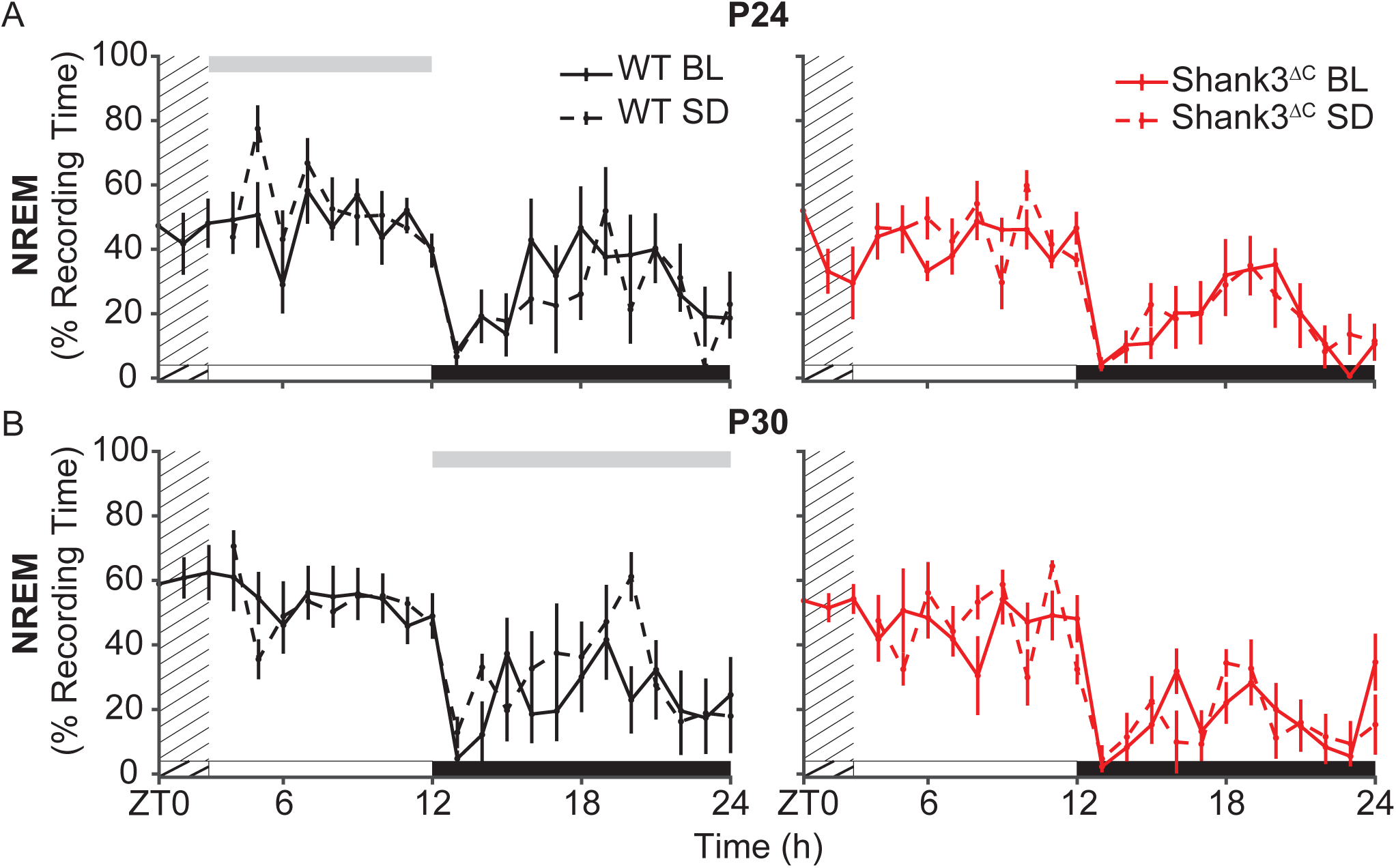
Shank3^ΔC^ animals do not increase sleep amounts following SD at P24 or P30. Total recording time (TRT) in NREM sleep during recovery sleep compared to baseline. **A**) P24. **B**) P30. The gray bar above the plots represents significance from repeated measures ANOVA across hour 3-12 (light period) and 13-24 (dark period) (p < 0.05). Light period is represented by the white bar, dark period by the black bar. The solid line represents the baseline day (BL) and the dashed line represents the recovery sleep day with the grated area representing 3 hours of SD. WT animals are represented in black (n = 5 P24, 6 P30) and Shank3^ΔC^ animals are represented in red (n = 6 P24, 6 P30).

Given that the inability to properly respond to SD is present in juvenile mutants, and that in adults we observe the largest effects of the mutation occur after SD (and not RS), we asked what the impact of the mutation was on genome-wide gene expression after SD at P24 and P30. Figure 6A shows the logFC after SD in P24 WT and Shank3^ΔC^ mice compared to allowed to sleep in their home cage (HC). Like adult animals, many DEGs are shared across genotypes (3029, shown in grey) including IEGs *Arc* and *Fos*. P24 WT animals have 4262 genotype-specific DEGs (shown in black) that include those related to neuronal growth and development (*Wnt9b, Gadd45b, Sirt2*), protein kinase activity (*Mtor, Camk2a*), transcription (*Egr2, Ep300*), and oxidative stress responses (*Atp12a*). P24 Shank3^ΔC^ animals have fewer genotype-specific DEGs than WT (1888, shown in red) that include ribosome and proteosome components (*Rps14, Rps18, Psmd11*). To identify genotype-specific DEGs that may mediate normal versus abnormal molecular responses to SD at P24, we intersected the lists of both upregulated and downregulated DEGs across genotypes. Figure 6B shows that WT animals upregulate 2065 unique genes while mutants upregulate 865 unique genes after SD (shown in green). Figure 6C shows a similar pattern in the downregulated genes (shown in purple), with 2197 DEGs unique to the WT after SD and only 1023 unique to the mutants. Together, these results indicate that the Shank3^ΔC^ mutation inhibits the molecular homeostatic response to SD at P24, as seen in adults. Lists of total DEGs for P24 animals can be found in Additional File 12 and lists of DEGs from the P24 intersections can be found in Additional File 13.

**Figure 6.**
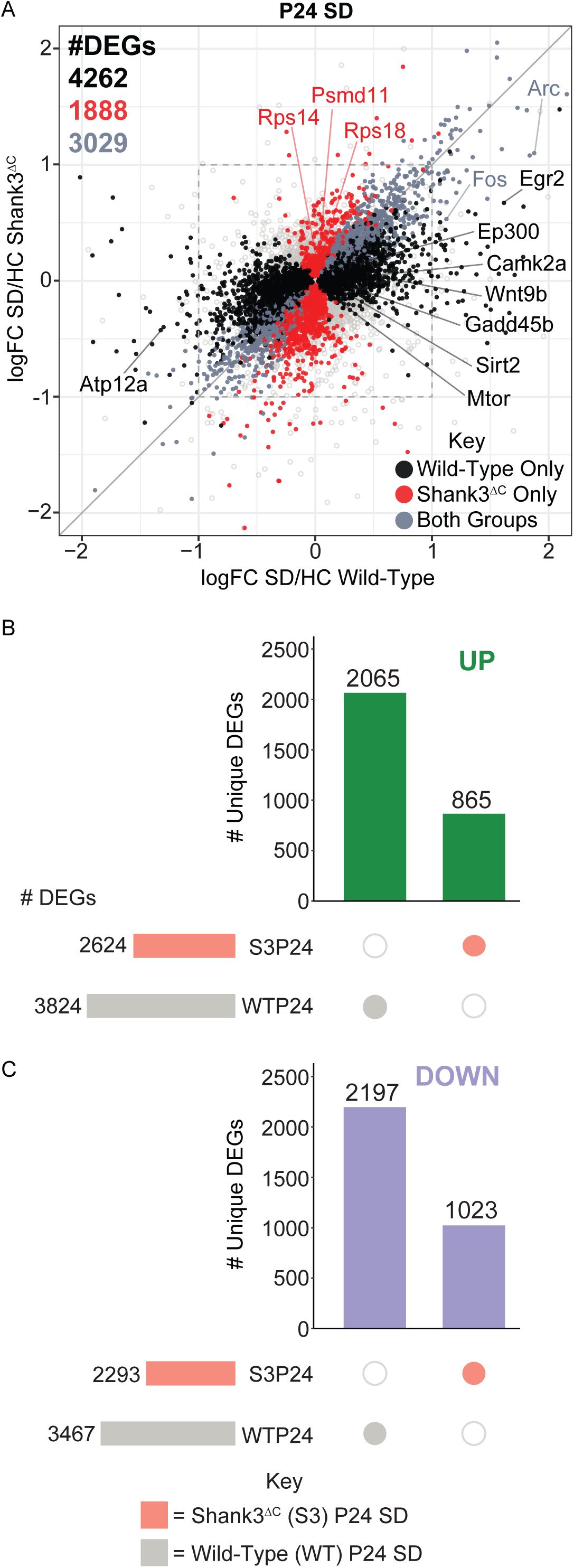
The Shank3^ΔC^ mutation impairs the typical transcriptional response to SD as early as P24. **A)** Correlation plot between the logFC of DEGs in P24 WT (x-axis) and Shank3^ΔC^ (y-axis) mice following SD/HC. DEGs found in both WT and Shank3^ΔC^ animals are shown in grey, DEGs unique to WT animals are in black and DEGs unique to Shank3^ΔC^ animals are in red. Plot windows are adjusted to show genes with a logFC ≤ |2|. The box represents a logFC cutoff of ± 1. Select genes of interest are highlighted. **B-C**) UpSet plots of the intersections of the number of DEGs between mutants (light red) and WT (light grey) across the P24 response to SD. Lists of DEGs were intersected for **B**) upregulated DEGs (green) and **C**) downregulated DEGs (purple). The total number of DEGs for each condition are shown in the set size rows on the left of the plot. The vertical bars represent the number of DEGs unique to the subset indicated by the colored dots in the intersection matrix. Intersections selected for functional enrichment analysis are shown.

After identifying genotype-specific DEGs following SD, we performed functional enrichment analysis to define functions that underlie the differences seen in the mutants’ response to SD at P24 (Figure 7). Figure 7A shows functions enriched in DEGs after SD unique to WT at P24. Upregulated functions form two clusters. Cluster one includes neurodevelopmental and signaling processes, such as neurogenesis and differentiation (*Mef2d, Wnt9b, Dvl1, Gadd45b),* whereas cluster two involves kinases and intracellular signaling pathways (*Camk2a, Mtor, Sik1).* Additional enriched upregulated functions include transcription regulation (*Ep300, Sirt2, Smarca4*, *Egr2*), axon guidance (*Camk2a*) and cell adhesion. Downregulated functions form two clusters, one of pathways involved in synaptic vesicle formation and oxidative phosphorylation (*Ndufa9*, *Cox5a/b*) and a second cluster of neurodegenerative disease-related pathways including Huntington’s, Parkinson’s, and Alzheimer’s disease (*Cox6a2*, *Ctnnb1*). These are similar to the pathways we see downregulated in adult WT animals. Additional downregulated functions include protein synthesis and DNA base excision repair. Figure 7B shows functions enriched in DEGs after SD unique to mutants at P24. Upregulated functions include protein synthesis and processing (*Rps18*, *Rps4x*, *Eif4a1/2*), neurodegenerative disease-related pathways (Alzheimer’s, Parkinson’s, Huntington’s) and processes related to receptor activity and glutamatergic synapses. Downregulated functions are associated with immune and inflammatory signaling as well as transcriptional regulation and DNA-binding (*Nfatc1, Dnmt1, Creb3, Tfap2a)*. Overall, while the Shank3^ΔC^ mutation does not substantially decrease the number of biological functions recruited after SD at P24, the mutants have an abnormal molecular response to SD when compared to WT that includes the upregulation of oxidative stress pathways linked to neurodegeneration and protein processing. The annotated results of functional enrichment analysis for Figure 8 can be found in Additional File 14. We previously showed that the ability to decrease sleep onset latency following SD develops between P24 and P30 in WT mice and this development is disrupted in Shank3^ΔC^ mutants (6); therefore, we next focused on defining the effect of the mutation on the molecular response to SD at P30. Surprisingly, we found that mutants have a larger transcriptional response to SD at P30 compared to WT (Figure 8). Figure 8A shows the logFC of all expressed genes in P30 WT and Shank3^ΔC^ mice after SD compared to allowed to sleep animals, with Shank3^ΔC^ animals having more genotype-specific DEGs (2453, shown in red) than WT (1497 DEGs, shown in black). Shank3^ΔC^-specific DEGs include initiation factors (*Eif4e*), unfolded protein response genes (*Hspa1a*), protein kinases (*Pik3r1*) and DNA repair genes (*Parpbp*), while WT-specific DEGs include developmental genes (*Pdgfb*) and transcription regulators *(Rora*). DEGs seen in both genotypes (2244, shown in grey) include IEGs (*Arc, Egr2*) and other elements of the unfolded protein response (*Hspa1b*). We further characterized P30 DEGs unique to WT or Shank3^ΔC^ by intersecting upregulated and downregulated DEG lists across genotypes. Figure 8B shows that WT animals upregulate 829 unique genes, while the Shank3^ΔC^ mutants upregulate 1149 unique genes after SD. Figure 8C shows that there are only 668 downregulated genes unique to the WT after SD but 1303 unique to the mutants. These results show that unlike at P24 or in adults, the Shank3^ΔC^ mutation produces a larger molecular homeostatic response to SD at P30. Lists of total DEGs in P30 animals are in Additional File 15 and lists of genotype-specific DEGs for P30 intersections are in Additional File 16. Additional File 17 shows data processing and normalization quality control plots (Additional File 17 A, B) and MA plots of differential expression for each of the P24 and P30 pairwise comparisons (Additional File 17 C-F).

**Figure 7.**
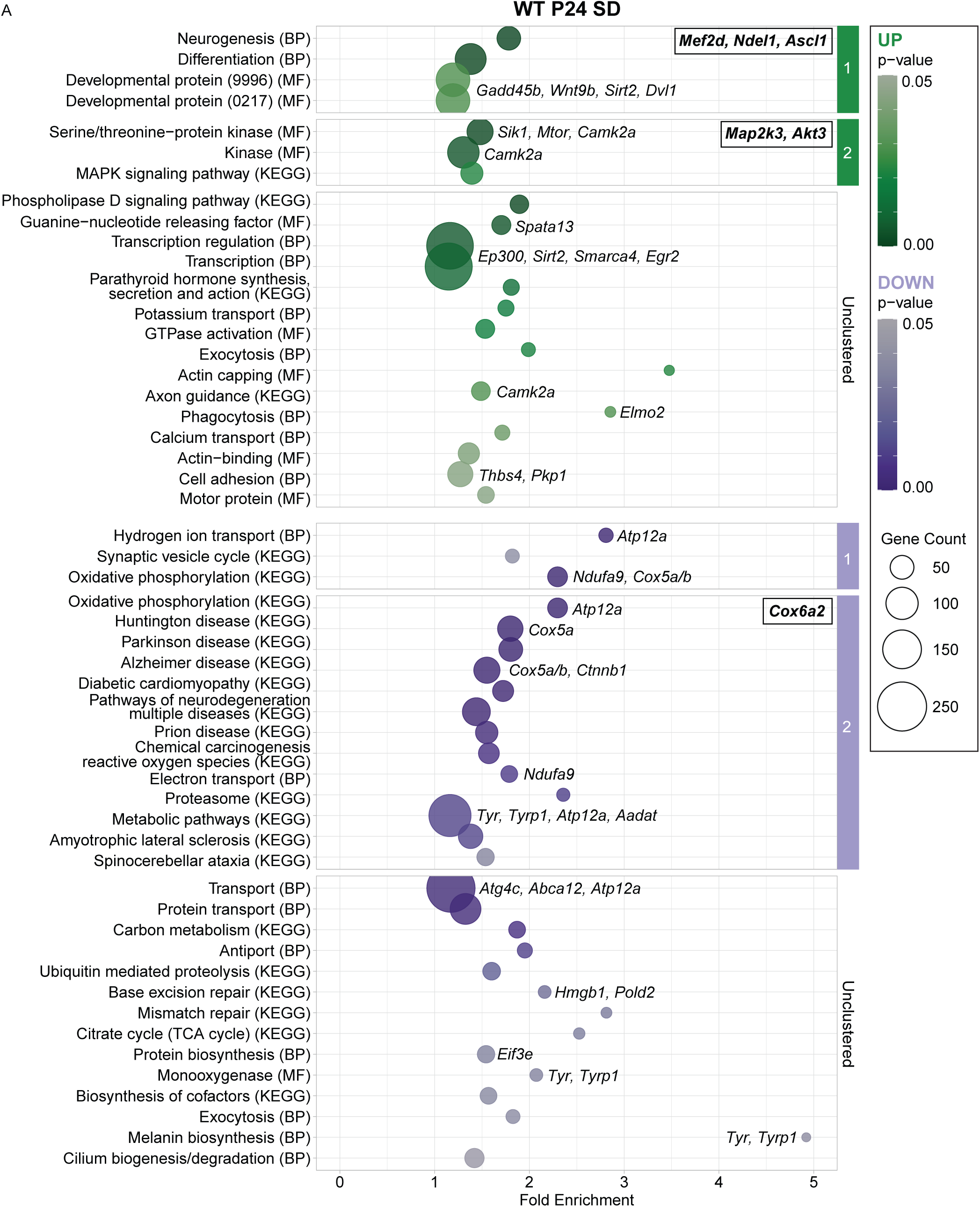
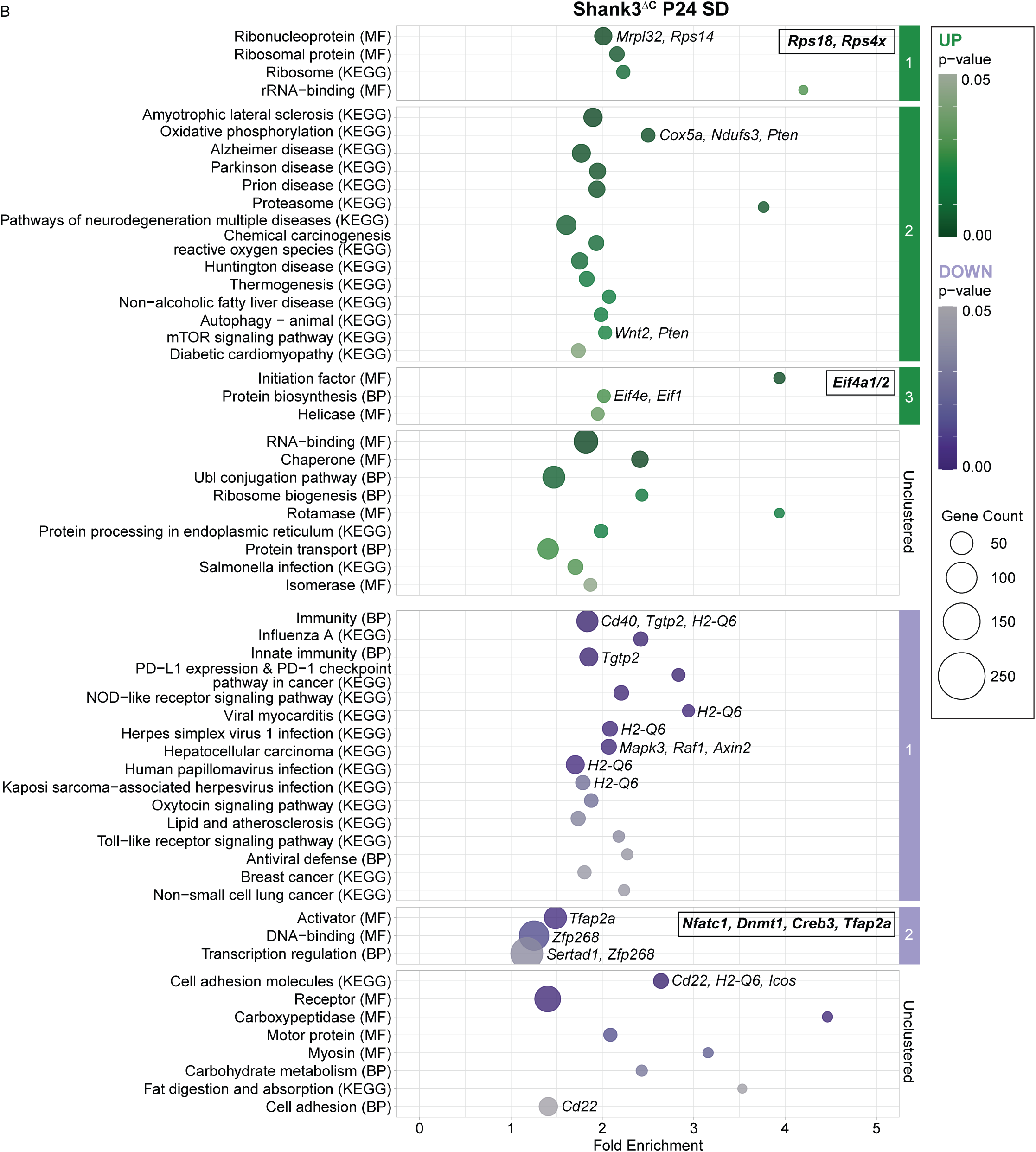
SD upregulates neurodevelopmental pathways at P24 in WT but not Shank3^ΔC^ animals. Functional enrichment analysis plots for DEGs unique to 3 hours of SD in P24 **A**) WT animals and **B**) Shank3^ΔC^ animals. Enriched functional annotation terms (modified Fisher’s exact *P* value < 0.05) from UniProt BP, MF and KEGG are displayed vertically. Circle size indicates the number of genes found within a given term which are plotted based on fold enrichment as displayed on the x-axis. Upregulated terms are in green and downregulated terms are in purple with darker shades representing smaller *P* values. Hub genes are bolded and displayed in boxes in the upper right corner of their corresponding cluster. Enrichment scores for each cluster in **A**: upregulated cluster 1 (2.58), upregulated cluster 2 (2.31), downregulated cluster 1 (3.28), downregulated cluster 2 (2.95). Enrichment scores for each cluster in **B**: upregulated cluster 1 (2.26), upregulated cluster 2 (2.16), upregulated cluster 3 (1.93), downregulated cluster 1 (2.15), downregulated cluster 2 (1.85).

**Figure 8.**
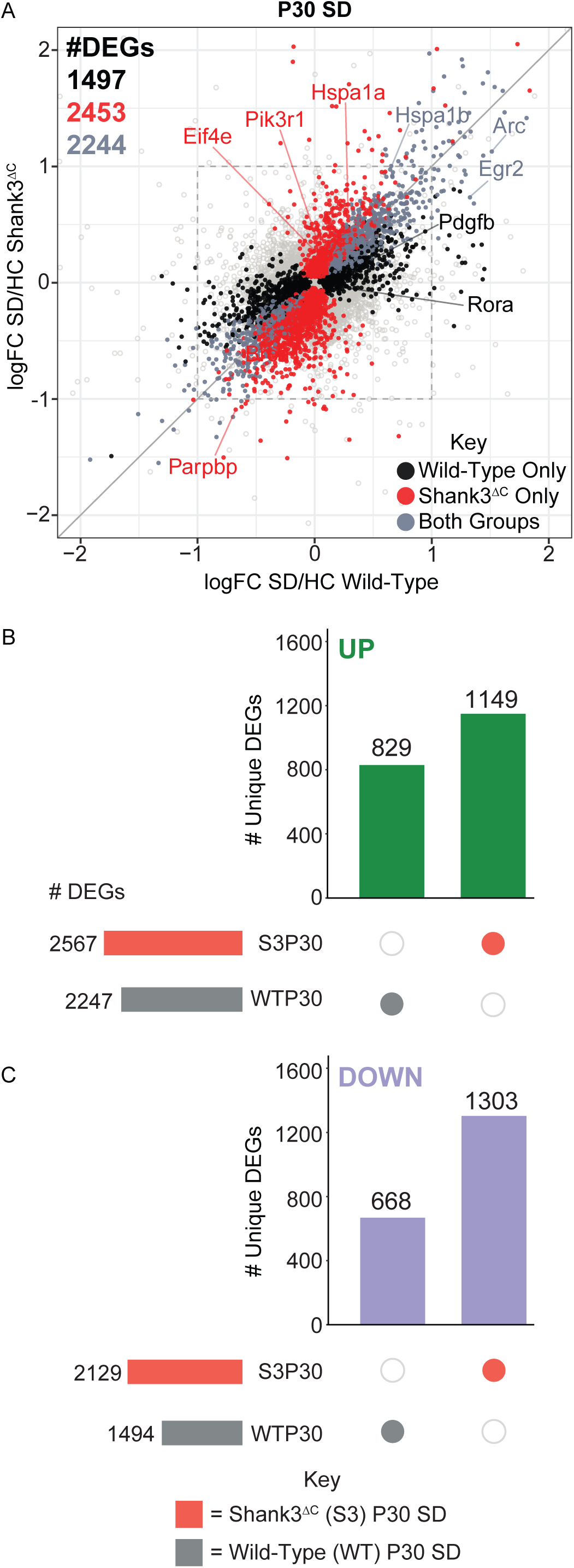
Shank3^ΔC^ animals have a larger response to SD at P30 than WT animals. **A**) Correlation plot between the logFC of DEGs in P30 WT (x-axis) and Shank3^ΔC^ (y-axis) mice following SD/HC. DEGs found in both WT and Shank3^ΔC^ animals are shown in grey, DEGs unique to WT animals are in black and DEGs unique to Shank3^ΔC^ animals are in red. Plot windows are adjusted to show genes with a logFC ≤ |2|. The box represents a logFC cutoff of ± 1. Select genes of interest are highlighted. **B-C**) UpSet plots of the intersections of the number of DEGs between Shank3^ΔC^ (light red) and WT (light grey) across the P30 response to SD. Lists of DEGs were intersected for **B**) upregulated DEGs (green) and **C**) downregulated DEGs (purple). The total number of DEGs for each condition are shown in the set size rows on the left of the plot. The vertical bars represent the number of DEGs unique to the subset indicated by the colored dots in the intersection matrix. Intersections selected for functional enrichment analysis are shown.

To identify functions and pathways that might underlie the enriched response to SD in P30 mutants, functional enrichment analysis was once again performed (Figure 9). Figure 9A shows functions enriched in DEGs after SD unique to WT at P30, which were fewer than at P24 and include upregulation of transcriptional activation and downregulation of cellular metabolism and growth (*Prkaca, Prcb1*). Figure 9B shows functions enriched in DEGs after SD unique to mutants at P30. These include upregulation of growth factors and oncogenic signaling (*Nras, Pik3r1, Hspa1a*) and downregulation of sugar metabolism, DNA damage/repair (*Parpbp*) as well as immune-related pathways. Overall, the mutation increases the biological functions recruited by SD in P30 mutants compared to WT but leads to a similar response to what is seen in P24 and adult Shank3^ΔC^ animals with the upregulation of stress-related pathways. Additional File 18 contains the annotated DAVID outputs for the P30 analysis.

**Figure 9.**
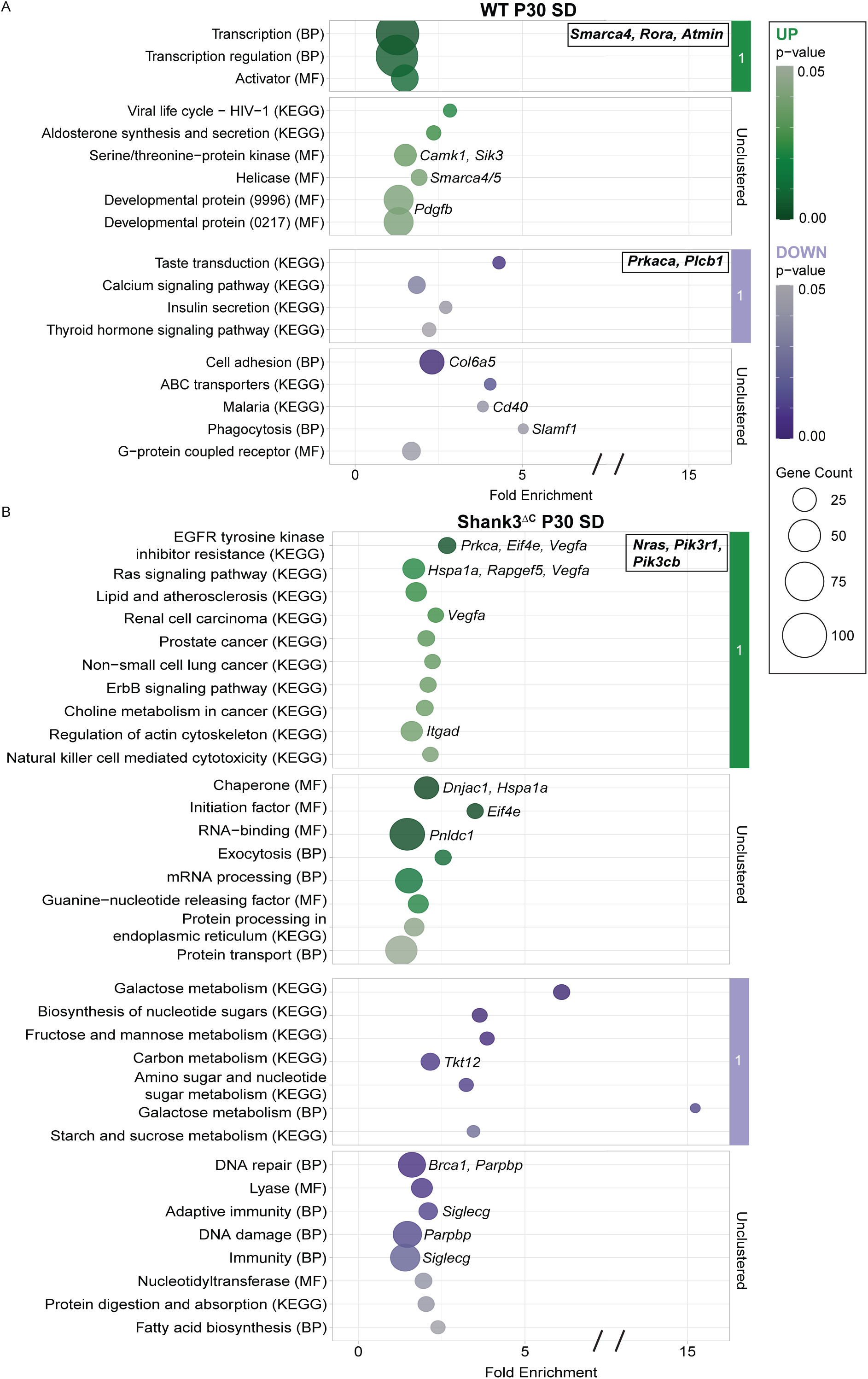
P30 Shank3^ΔC^ animals upregulate protein synthesis and downregulate metabolic pathways following SD. Functional enrichment analysis plots for DEGs unique to 3 hours of sleep SD in P30 **A)** WT animals and **B**) Shank3^ΔC^ animals. Enriched functional annotation terms (modified Fisher’s exact *P* value < 0.05) from UniProt BP, MF and KEGG are displayed vertically. Circle size indicates the number of genes found within a given term which are plotted based on fold enrichment as displayed on the x-axis. Upregulated terms are in green and downregulated terms are in purple with darker shades representing smaller *P* values. Hub genes are bolded and displayed in boxes in the upper right corner of their corresponding cluster. Enrichment scores for each cluster in **A**: upregulated cluster 1 (2.29), downregulated cluster 1 (1.61). Enrichment scores for each cluster in **B**: upregulated cluster 1 (1.61), downregulated cluster 1 (2.56).

### The Shank3^ΔC^ mutation engages neurodegenerative pathways after SD regardless of age

Although there are clear differences in the transcriptional response to SD and the effect of the Shank3^ΔC^ mutation across ages, we were interested in identifying functions that were common across ages and may mediate sleep homeostasis deficits in the mutants regardless of age. We integrated functional enrichment analyses from Figure 3 (adult), Figure 7 (P24) and Figure 9 (P30) to find pathways that were common across genotypes and ages following SD (Figure 10). Figure 10 shows functions that are enriched in at least three groups and contain at least 20 genes with upregulated functions in green and downregulated functions in purple. Neurodegenerative pathways (e.g. Alzheimer’s, Huntington’s and Parkinson’s) are consistently downregulated in WT but upregulated in mutants across ages. Additionally, protein transport pathways and ribosomal components are downregulated in WT but upregulated in mutants. Transcription is upregulated in both WT and Shank3^ΔC^ mutants, indicating that the difference between the genotypes is not related to transcriptional regulation alone. Overall, this integration shows that SD has the opposite effect on oxidative stress seen in neurodegenerative disease-related pathways and protein synthesis in WT and Shank3^ΔC^ mutants at all ages.

**Figure 10.**
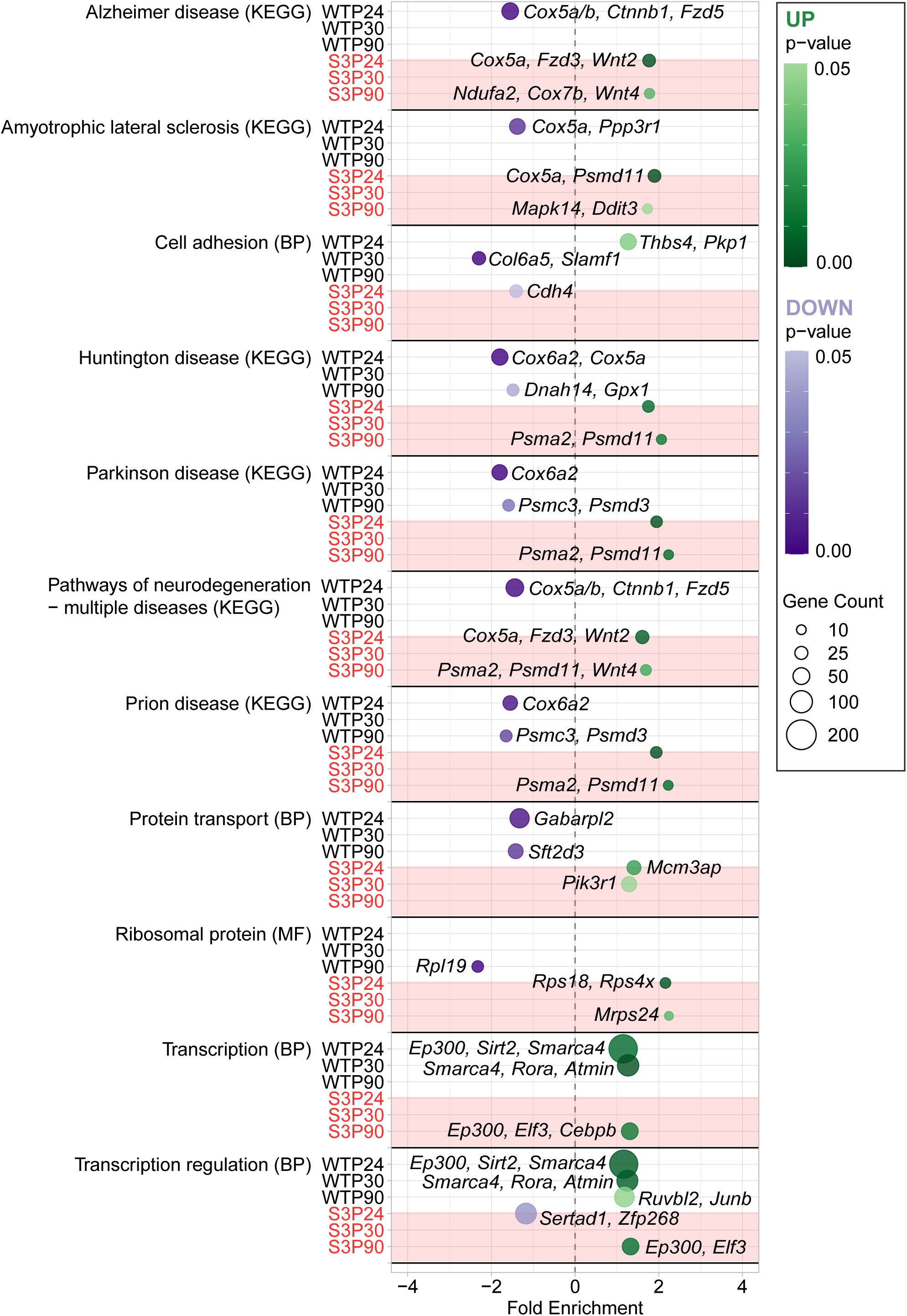
Pathways related to oxidative stress are downregulated in WT but upregulated in Shank3^ΔC^ across ages. Functional enrichment analysis plot for pathways unique to SD but shared across ages and genotypes. Enriched functional annotation terms (modified Fisher’s exact *P* value < 0.05) from UniProt BP, MF and KEGG are displayed vertically. Circle size indicates the number of genes found within a given term and which are plotted based on fold enrichment as displayed on the x-axis. Upregulated terms are in green and downregulated terms are in purple with darker shades representing smaller *P* values. Terms where at least one condition has a gene count ≥ 20 are displayed.

## Discussion

In this study, we examined the impact of the Shank3^ΔC^ mutation on the genome-wide cortical gene expression response to acute SD from juveniles to adulthood in male mice. Comparing gene expression across age and genotype allowed for the identification of condition-specific responses to SD and subsequent recovery sleep (SD+RS), helping us understand the potential molecular basis of sleep problems in Shank3^ΔC^ mutants. In adults, the mutation substantially lowers the number of DEGs that respond to SD, reduces the ability of RS to recover gene expression changes and decreases the number of enriched biological pathways and functions recruited following SD and SD+RS (Figures 1,2,3,4). As sleep homeostasis deficits are observed not only in adult mutants but also at P24 and P30 (Figure 5), we next investigated the impact of the mutation on the molecular response to SD at these ages. Mutants have a smaller transcriptional response to SD at P24 but a larger response at P30 when compared to WT (Figures 6,8). While P24 mutants and WT have a similar number of enriched functions after SD, at P30 mutants have more enriched pathways than age-matched WT animals (Figure 7,9). Integrating the results of functional enrichment analysis across ages and genotypes shows that mutants consistently upregulate oxidative stress pathways related to neurodegeneration and protein synthesis regardless of age while WT downregulate these functions (Figure 10). Together, these findings indicate that the Shank3^ΔC^ mutation interferes with the ability to respond properly to SD at the molecular level during development.

Our data show that the effect of the Shank3^ΔC^ mutation on gene expression after acute SD is different at P24 than at P30. WT animals have a decrease in the number of DEGs following SD as we previously described (17), while Shank3^ΔC^ mutants have a similar number of DEGs at both ages (Figure 6,8). It is known that, in rodents, sleep homeostasis matures postnatally between P12 and P30 (6,27,28). It is possible that the reduction in gene expression level response to SD in WT reflects an increased ability to stay awake as animals get older. In other words, 3 hours of SD at P24 produces a proportionally larger increase in sleep pressure compared to the later developmental age of P30 as our previous work suggests (6). Shank3^ΔC^ mutants show a similar number of DEGs between P24 and P30 (4917 vs 4697) and display a substantial reduction in the overlap with WT DEGs after SD (3029 vs 2244) (Figure 6,8). We know that at the physiological level the Shank3^ΔC^ response to SD at P30 is similar to the P24 response as evidenced by their inability to fall asleep fast when sleepy, while in WT it has matured (6). Our findings suggest that this is also true at the molecular level. Thus, the Shank3^ΔC^ mutation may prevent mutants from developing typical homeostatic sleep regulation between P24 and P30. Our data provides insight into what the molecular mechanisms underlying this typical maturation may be. P24 and P30 WT (but not mutant) animals upregulate neurodevelopmental pathways and developmental proteins in response to SD (Figures 7,9), although this genotype specific difference is proportionally smaller in P30 animals. In adult WT (but not mutant animals) pathways recruited by SD now include long-term potentiation, intracellular signaling and DNA damage/repair (Figure 3). These findings align with previous work showing that the typical response to SD in adult WTs involves the induction of IEGs associated with synaptic plasticity and DNA damage/repair (7,12,16,17). Overall, in WT animals, SD alters pathways associated with neuronal activity (as expected from extended wake) shifting from neurodevelopmental pathways in juveniles to LTP and DNA damage/repair in adults. The fact that Shank3^ΔC^ mutants are unable to upregulate these pathways could indicate they cannot respond properly to increased neuronal activity that may drive the sleep homeostat at the molecular/cellular level.

Our data also show that mutants produce and aberrant response to SD regardless of age. Shank3^ΔC^ mutants upregulate pathways related to neurodegenerative diseases (Alzheimer’s, Huntington’s, Parkinson’s, etc.) as well as molecular functions involved in ribosome biogenesis (protein processing, ribosomal proteins, etc.) following SD, indicating an increase in oxidative stress in the brain (Figures 3, 7, 9, 10). This is particularly interesting given that WT downregulate these processes in our study. Extended wakefulness is associated with increased production of reactive oxidative species (ROS) in neurons, as active neurons during wake require greater energy expenditure and therefore greater mitochondrial ATP production (30,31). The downregulation of these pathways in WT animals can be interpreted as an attempt to mitigate increased cellular stress induced by SD (Figures 3, 7, 9, 10). Thus, our data suggests that in addition to not being able to engage neuronal activity-dependent pathways, Shank3^ΔC^ mutants, by upregulating energy intensive pathways involved in oxidative phosphorylation and protein synthesis, may worsen the adverse effects of acute SD through an increase in oxidative stress.

### Limitations

While our findings provide new insight into the transcriptional response to SD across development and in Shank3^ΔC^ mice, our study also contained some limitations. Only male mice were used for this research, allowing for a more direct comparison of previous work but preventing us from analyzing the female transcriptional response to SD, which may differ given differences in the timeline of brain maturations between males and females (32,33). Additionally, this study utilized a single rodent model of ASD – Shank3^ΔC^ mutants. Whether it is true that other genetic mutations associated with ASD may worsen the homeostatic response to SD at the molecular level remains to be explored. Our previous work using Mecp2^-/y^ animals suggests this is true at the physiological level (34); thus, further studies using other mouse models are warranted. Last, our findings cannot provide conclusions for the effect of SD or RS on other brain regions or specific cell types, which is an important consideration as it has been previously shown that SD primarily impacts excitatory neuronal cells (15). Future work using single-cell and spatial transcriptomic technologies across brain regions will be key to understanding the effects of SD and its interaction with ASD-linked mutations at the cellular level.

### Conclusions

In this study, we showed that the Shank3^ΔC^ mutation impairs the neurotypical gene expression response to acute sleep deprivation in juvenile and adult male mice. We found that the mutation largely represses the transcriptional response to SD, leads to the consistent upregulation of oxidative stress and neurodegeneration, and prevents the upregulation of processes such as neurodevelopment and DNA damage repair, which differ across ages. We conclude that the effect of the Shank3^ΔC^ mutation at the molecular level is an impairment in how animals track and respond to sleepiness. Our findings suggest that enhancing the recruitment of neurotypical pathways related to sleep homeostasis (neuronal growth at young ages or DNA repair pathways in adulthood) while diminishing the impact of oxidative stress following SD may be actionable strategies to improve sleep in individuals with ASD. Future work will focus on replicating these findings across different ASD animal models, determining cell-type and circuit specificity as well as validating the involvement of prioritized pathways.

## List of abbreviations

ASD: Autism Spectrum Disorder
WT: wild-type
P24: postnatal day 24
P30: postnatal day 30
P90: postnatal day 90
SD: sleep deprivation
SD+RS: sleep deprivation and recovery sleep
HC: home cage
ZT: Zeitgeber time
NREM: non-rapid eye movement sleep
REM: rapid eye movement sleep
DEG: differentially expressed genes
MA: minus-average
IEG: immediate early genes
GEO: Gene Expression Omnibus Database
ROS: reactive oxidative species
DAVID: Database for Annotation, Visualization, and Integrated Discovery
BP: biological processes
MF: molecular function
KEGG: Kyoto Encyclopedia of Genes and Genomes

## Declarations Ethics declaration

All experimental procedures were approved by the Institutional Care and Use Committee of Washington State University and conducted in accordance with National Research Council guidelines and regulations for experiments in live animals.

## Consent for publication

Not applicable

## Availability of data and materials

Sequencing data are available through the Gene Expression Omnibus Database (GEO) under accession number GSE306621. Data from wild-type animals used in this study were previously published in GSE211301 (Muheim et al. 2023, SD and HC P24, P30) and GSE113754 (Ingiosi et al. 2019, SD and HC adult). EEG data are available at https://sleepdata.org/datasets/medina-2022. All code used for analysis and visualization is available on Github: https://github.com/PeixotoLab/RNAseq_sleep_developmentShank3

## Authors’ contributions

Concept: L.P.

Design: E.M., C.M., K.S., L.P.

Writing: E.W., L.P., E.M., C.O. K.S.

Collection: E.M., C.M., K.S., K.F., T.W.P., A.I, L.P

Analysis: E.W., C.O, L.P.

## Acknowledgements

Not Applicable

## Trial registration

Not Applicable

## Additional Files

Supplement to Figure 1:

- Additional File 1; P90 total DEGs; .xlsx; lists of total DEGs in P90 animals

- Additional File 2; P90 WT and Shank3^ΔC^ mice have unique gene expression profiles following SD and SD+RS; .docx; quality control plots showing the results of normalization and differential expression analysis for P90 animals

Supplement to Figure 2:

- Additional File 3; P90 unique DEGs; .xlsx; lists of unique DEGs obtained by intersecting total DEG lists from P90 animals

Supplement to Figure 3:

- Additional File 4; annotated DAVID P90 SD; .xlsx; annotated results of functional enrichment analysis for P90 SD samples

Supplement to Figure 4:

- Additional File 5; annotated DAVID P90 RS; .xlsx; annotated results of functional enrichment analysis for P90 RS samples

Supplement to Figures 1-4:

- Additional File 6; a subset of genes is differentially expressed following SD, SD+RS regardless of genotype at P90; .docx; upset plots showing DEGs that are common across multiple conditions in P90 animals

- Additional File 7; P90 common DEGs; .xlsx; lists of DEGs that are common across multiple conditions in P90 animals

- Additional File 8; Sleep manipulations impact DNA damage detection, circadian rhythms, stress responses regardless of genotype at P90; .docx; functional enrichment analysis plots for common intersections in P90 samples

- Additional File 9; annotated DAVID P90 common; .xlsx; annotated results of functional enrichment analysis for genes in the common P90 intersections

Supplement to Figure 5:

- Additional File 10; WT animals are awake less following SD at P30 while Shank3^ΔC^ animals show differences in REM sleep; .docx; time in state traces for wake and REM sleep in P24 and P30 animals

- Additional File 11; P24 P30 EEG statistics; .xlsx; statistics for time in state analysis in P24 and P30 animals

Supplement to Figure 6:

- Additional File 12; P24 total DEGs; .xlsx; lists of total DEGs for P24 samples

- Additional File 13; P24 unique DEGs; .xlsx; lists of unique DEGs obtained by intersecting total DEG lists of P24 animals

Supplement to Figure 7:

- Additional File 14; annotated DAVID P24; .xlsx; annotated results of functional enrichment analysis for P24 samples

Supplement to Figure 8:

- Additional File 15; P30 total DEGs; .xlsx; lists of total DEGs for P30 samples

- Additional File 16; P30 unique DEGs; .xlsx; lists of unique DEGs obtained by intersecting total DEG lists of P30 animals

Supplement to Figures 6, 8:

- Additional File 17; WT and Shank3^ΔC^ gene expression differs following SD at P24 and P30; .docx; quality control plots showing the results of normalization and differential expression analysis for P24/P30 animals

Supplement to Figure 9:

- Additional File 18; annotated DAVID P30; .xlsx; annotated results of functional enrichment analysis for P30 samples

## References

1. Petruzzelli MG, Matera E, Giambersio D, Marzulli L, Gabellone A, Legrottaglie AR, et al. Subjective and Electroencephalographic Sleep Parameters in Children and Adolescents with Autism Spectrum Disorder: A Systematic Review. J Clin Med. 2021 Aug 30;10(17):3893. doi:10.3390/jcm10173893 PubMed PMID: 34501341; PubMed Central PMCID: PMC8432113.

2. MacDuffie KE, Shen MD, Dager SR, Styner MA, Kim SH, Paterson S, et al. Sleep Onset Problems and Subcortical Development in Infants Later Diagnosed With Autism Spectrum Disorder. AJP. 2020 May 7;177(6):518–25. doi:10.1176/appi.ajp.2019.19060666

3. Hacohen M, Levy A, Kaiser H, Green Snyder L, Amatya A, Gundersen BB, et al. An open science resource for accelerating scalable digital health research in autism and other neurodevelopmental conditions. Nat Neurosci. 2026 Feb;29(2):467–78. doi:10.1038/s41593-025-02146-3

4. Ingiosi A, Schoch H, Wintler TP, Singletary KG, Righelli D, Roser L, et al. Shank3 Modulates Sleep and Expression of Circadian Transcription Factors. Kim E, editor. eLife. 2019 Apr 11;8:e42819. doi:10.7554/eLife.42819

5. Medina E, Rempe MJ, Muheim C, Schoch H, Singletary K, Ford K, et al. Sex differences in sleep deficits in mice with an autism-linked Shank3 mutation. Biol Sex Differ. 2024 Oct 28;15(1):85. doi:10.1186/s13293-024-00664-6 PubMed PMID: 39468684; PubMed Central PMCID: PMC11514800.

6. Medina E, Schoch H, Ford K, Wintler T, Singletary KG, Peixoto L. Shank3 influences mammalian sleep development. Journal of Neuroscience Research. 2022;100(12):2174–86. doi:10.1002/jnr.25119

7. Maret S, Dorsaz S, Gurcel L, Pradervand S, Petit B, Pfister C, et al. Homer1a is a core brain molecular correlate of sleep loss. Proc Natl Acad Sci U S A. 2007 Dec 11;104(50):20090–5. doi:10.1073/pnas.0710131104 PubMed PMID: 18077435; PubMed Central PMCID: PMC2148427.

8. Wisor JP, Pasumarthi RK, Gerashchenko D, Thompson CL, Pathak S, Sancar A, et al. Sleep Deprivation Effects on Circadian Clock Gene Expression in the Cerebral Cortex Parallel Electroencephalographic Differences among Mouse Strains. J Neurosci. 2008 Jul 9;28(28):7193–201. doi:10.1523/JNEUROSCI.1150-08.2008 PubMed PMID: 18614689.

9. Gerstner JR, Koberstein JN, Watson AJ, Zapero N, Risso D, Speed TP, et al. Removal of unwanted variation reveals novel patterns of gene expression linked to sleep homeostasis in murine cortex. BMC Genomics. 2016 25;17(Suppl 8):727. doi:10.1186/s12864-016-3065-8 PubMed PMID: 27801296; PubMed Central PMCID: PMC5088519.

10. Naidoo N, Giang W, Galante RJ, Pack AI. Sleep deprivation induces the unfolded protein response in mouse cerebral cortex. Journal of Neurochemistry. 2005 Feb 4. 10.1111/j.1471-4159.2004.02952.x

11. Mackiewicz M, Shockley KR, Romer MA, Galante RJ, Zimmerman JE, Naidoo N, et al. Macromolecule biosynthesis: a key function of sleep. Physiol Genomics. 2007 Nov 14;31(3):441–57. doi:10.1152/physiolgenomics.00275.2006 PubMed PMID: 17698924.

12. Cirelli C, Gutierrez CM, Tononi G. Extensive and Divergent Effects of Sleep and Wakefulness on Brain Gene Expression. Neuron. 2004 Jan 8;41(1):35–43. doi:10.1016/S0896-6273(03)00814-6

13. Vecsey CG, Peixoto L, Choi JHK, Wimmer M, Jaganath D, Hernandez PJ, et al. Genomic analysis of sleep deprivation reveals translational regulation in the hippocampus. Physiol Genomics. 2012 Oct 17;44(20):981–91. doi:10.1152/physiolgenomics.00084.2012 PubMed PMID: 22930738; PubMed Central PMCID: PMC3472468.

14. Tudor JC, Davis EJ, Peixoto L, Wimmer ME, van Tilborg E, Park AJ, et al. Sleep deprivation impairs memory by attenuating mTORC1-dependent protein synthesis. Sci Signal. 2016 Apr 26;9(425):ra41. doi:10.1126/scisignal.aad4949 PubMed PMID: 27117251; PubMed Central PMCID: PMC4890572.

15. Ford K, Zuin E, Righelli D, Medina E, Schoch H, Singletary K, et al. A global transcriptional atlas of the effect of acute sleep deprivation in the mouse frontal cortex. iScience. 2024 Sep 20;27(9). doi:10.1016/j.isci.2024.110752

16. Popescu A, Ottaway C, Ford K, Medina E, Wintler T, Ingiosi A, et al. Transcriptional Dynamics of Sleep Deprivation and Subsequent Recovery Sleep in the Male Mouse Cortex. Physiol Genomics. 2025 May 2. doi:10.1152/physiolgenomics.00128.2024 PubMed PMID: 40315180.

17. Muheim CM, Ford K, Medina E, Singletary K, Peixoto L, Frank MG. Ontogenesis of the molecular response to sleep loss. Neurobiol Sleep Circadian Rhythms. 2023 Mar 16;14:100092. doi:10.1016/j.nbscr.2023.100092 PubMed PMID: 37020466; PubMed Central PMCID: PMC10068260.

18. Patro R, Duggal G, Love MI, Irizarry RA, Kingsford C. Salmon provides fast and bias-aware quantification of transcript expression. Nat Methods. 2017 Apr;14(4):417–9. doi:10.1038/nmeth.4197 PubMed PMID: 28263959; PubMed Central PMCID: PMC5600148.

19. Love MI, Soneson C, Hickey PF, Johnson LK, Pierce NT, Shepherd L, et al. Tximeta: Reference sequence checksums for provenance identification in RNA-seq. PLoS Comput Biol. 2020 Feb;16(2):e1007664. doi:10.1371/journal.pcbi.1007664 PubMed PMID: 32097405; PubMed Central PMCID: PMC7059966.

20. Risso D, Schwartz K, Sherlock G, Dudoit S. GC-content normalization for RNA-Seq data. BMC Bioinformatics. 2011 Dec 17;12:480. doi:10.1186/1471-2105-12-480 PubMed PMID: 22177264; PubMed Central PMCID: PMC3315510.

21. Risso D, Ngai J, Speed TP, Dudoit S. Normalization of RNA-seq data using factor analysis of control genes or samples. Nat Biotechnol. 2014 Sep;32(9):896–902. doi:10.1038/nbt.2931 PubMed PMID: 25150836; PubMed Central PMCID: PMC4404308.

22. Zhu A, Srivastava A, Ibrahim JG, Patro R, Love MI. Nonparametric expression analysis using inferential replicate counts. Nucleic Acids Res. 2019 Oct 10;47(18):e105. doi:10.1093/nar/gkz622 PubMed PMID: 31372651; PubMed Central PMCID: PMC6765120.

23. Dyer SC, Austine-Orimoloye O, Azov AG, Barba M, Barnes I, Barrera-Enriquez VP, et al. Ensembl 2025. Nucleic Acids Res. 2025 Jan 6;53(D1):D948–57. doi:10.1093/nar/gkae1071

24. Sherman BT, Hao M, Qiu J, Jiao X, Baseler MW, Lane HC, et al. DAVID: a web server for functional enrichment analysis and functional annotation of gene lists (2021 update). Nucleic Acids Res. 2022 Jul 5;50(W1):W216–21. doi:10.1093/nar/gkac194 PubMed PMID: 35325185; PubMed Central PMCID: PMC9252805.

25. Frank MG, Heller HC. Development of REM and slow wave sleep in the rat. American Journal of Physiology-Regulatory, Integrative and Comparative Physiology. 1997 Jun 1;272(6):R1792–9. doi:10.1152/ajpregu.1997.272.6.R1792

26. Frank MG, Heller HC. Development of diurnal organization of EEG slow-wave activity and slow-wave sleep in the rat. American Journal of Physiology-Regulatory, Integrative and Comparative Physiology. 1997 Aug 1;273(2):R472–8. doi:10.1152/ajpregu.1997.273.2.R472

27. Frank MG, Morrissette R, Heller HC. Effects of sleep deprivation in neonatal rats. American Journal of Physiology-Regulatory, Integrative and Comparative Physiology. 1998 Jul 1. Located at: Bethesda, MD. doi:10.1152/ajpregu.1998.275.1.R148

28. Frank MG, Ruby NF, Heller HC, Franken P. Development of Circadian Sleep Regulation in the Rat: A Longitudinal Study Under Constant Conditions. Sleep. 2017 Mar 1;40(3):zsw077. doi:10.1093/sleep/zsw077

29. Dijk DJ, Duffy JF, Silva EJ, Shanahan TL, Boivin DB, Czeisler CA. Amplitude Reduction and Phase Shifts of Melatonin, Cortisol and Other Circadian Rhythms after a Gradual Advance of Sleep and Light Exposure in Humans. PLoS One. 2012 Feb 17;7(2):e30037. doi:10.1371/journal.pone.0030037 PubMed PMID: 22363414; PubMed Central PMCID: PMC3281823.

30. Hartmann C, Kempf A. Mitochondrial control of sleep. Current Opinion in Neurobiology. 2023 Aug 1;81:102733. doi:10.1016/j.conb.2023.102733

31. Wang L, Aton SJ. Perspective – ultrastructural analyses reflect the effects of sleep and sleep loss on neuronal cell biology. Sleep. 2022 May 1;45(5):zsac047. doi:10.1093/sleep/zsac047

32. Qiu LR, Fernandes DJ, Szulc-Lerch KU, Dazai J, Nieman BJ, Turnbull DH, et al. Mouse MRI shows brain areas relatively larger in males emerge before those larger in females. Nat Commun. 2018 Jul 5;9:2615. doi:10.1038/s41467-018-04921-2 PubMed PMID: 29976930; PubMed Central PMCID: PMC6033927.

33. Mannino GS, Green TRF, Murphy SM, Donohue KD, Opp MR, Rowe RK. The importance of including both sexes in preclinical sleep studies and analyses. Sci Rep. 2024 Oct 15;14(1):23622. doi:10.1038/s41598-024-70996-1

34. Al Maghribi A, Ottaway C, Rempe M, Medina E, Ford K, Singletary K, et al. Loss of MeCP2 leads to sleep deficits that are time-of-day dependent and worsen with sleep deprivation. Neurobiology of Sleep and Circadian Rhythms. 2025 Nov 1;19:100132. doi:10.1016/j.nbscr.2025.100132

