## Additional File 2 for "Shank3 mutation disrupts the molecular signature of sleepiness across development"

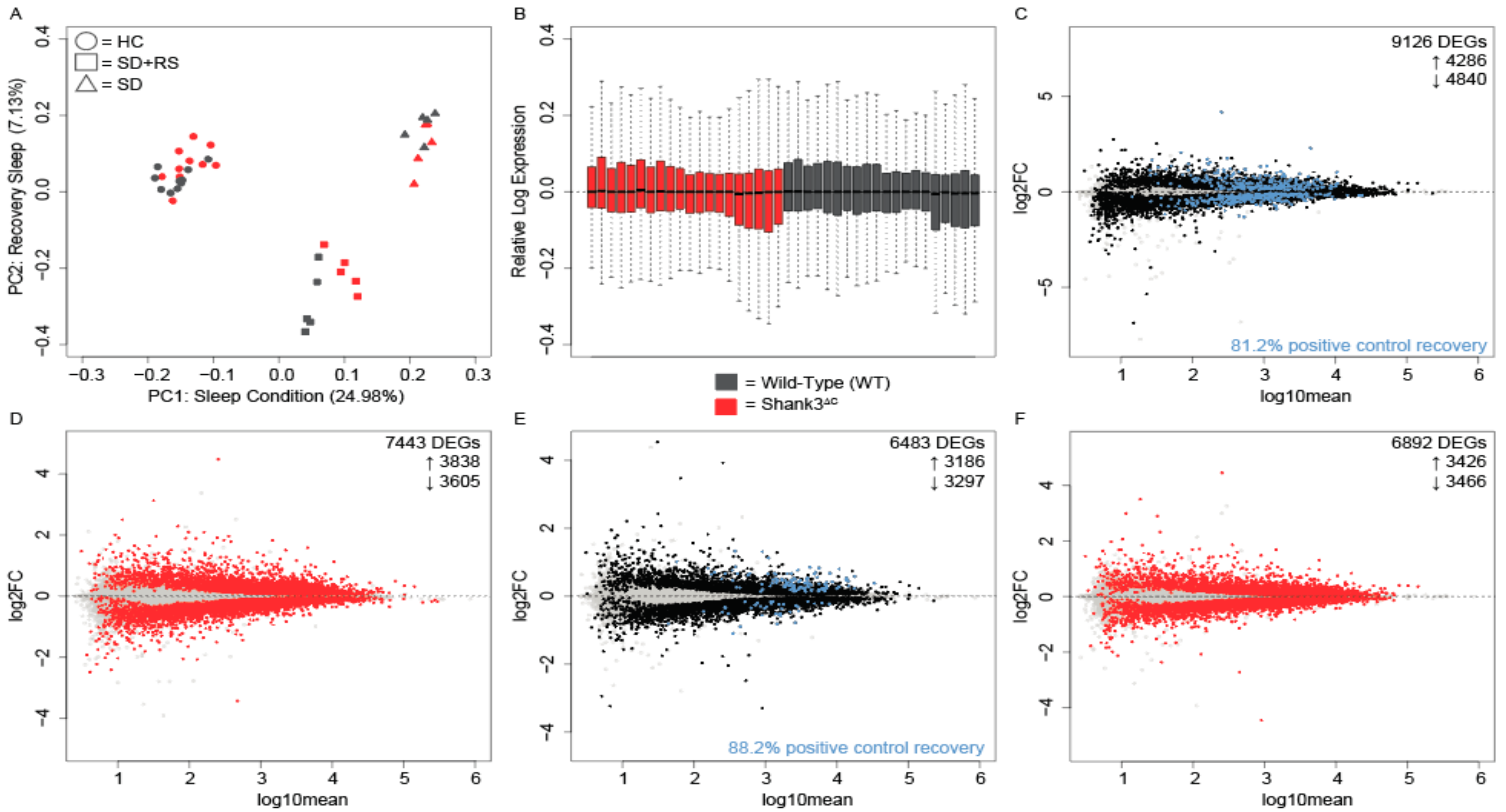


**Additional File 2**. **P90 WT and Shank3$^{\Delta C}$ mice have unique gene expression profiles following SD and SD+RS. A**) Principal component analysis following RUVs normalization with k = 15 unwanted factors. Circles represent home cage (HC) animals, squares represent animals that were sleep deprived for 5 hours then given a 2-hour recovery period (SD+RS) and triangles represent animals that were sleep deprived for 5 hours (SD). WT animals are shown in black and Shank3$^{\Delta C}$ animals are shown in red. n = 5 for each condition. Sleep condition (PC1, 24.98%) and recovery sleep (PC2, 7.13%) account for the most variability within the data. **B**) Relative log expression for each sample and condition. Color code as in A. **C-F**) MA plots following differential expression analysis on RUV normalized gene counts. **C**) WT SD, **D**) Shank3$^{\Delta C}$ SD, **E**) WT SD+RS, **F**) Shank3$^{\Delta C}$ SD+RS. Color code as in A showing differentially expressed genes (DEGs) in black for WT animals and red for Shank3$^{\Delta C}$ animals. Positive control genes are highlighted in blue for WT animals. Numbers of total DEGs, upregulated DEGs and downregulated DEGs are reported in the top right corner for each condition. Additionally, the percentage of recovered positive control genes are reported for the WT animals.
