## Additional File 6 for "Shank3 mutation disrupts the molecular signature of sleepiness across development"

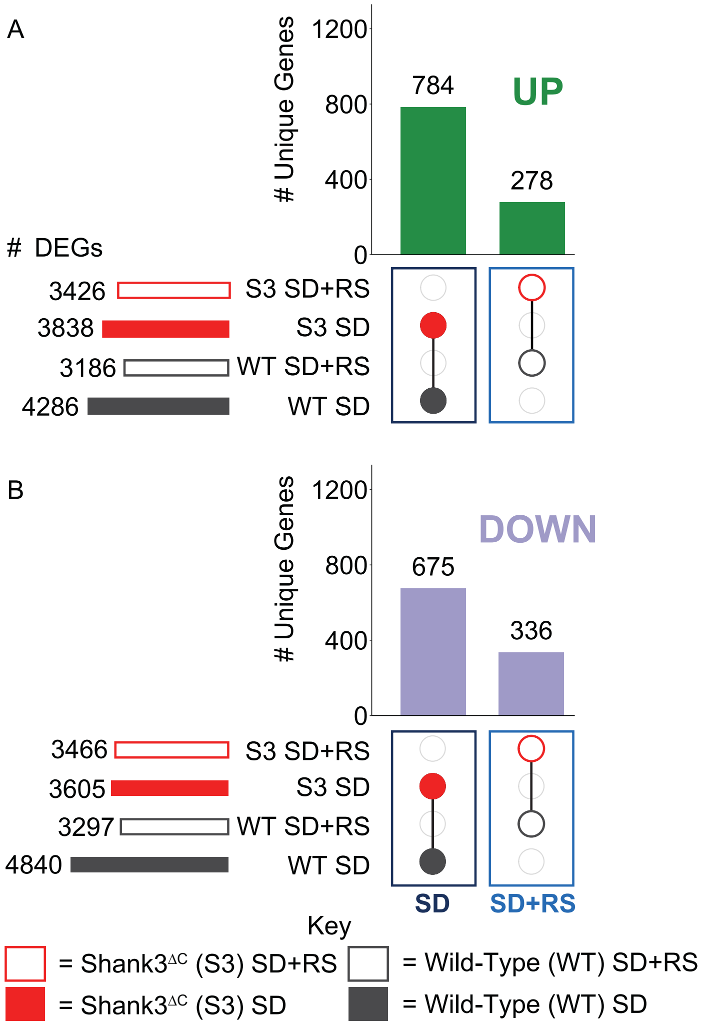


**Additional File 6. A subset of genes is differentially expressed following SD, SD+RS regardless of genotype in adults.** UpSet plots of the intersections of the number of DEGs between adult Shank3$^{\Delta C}$ (red) and WT (dark grey) response to SD (filled dots, navy blue box) or SD+RS (outlined dots, blue box). Lists of DEGs were intersected for **A**) upregulated DEGs (green) and **B**) downregulated DEGs (purple). The total number of DEGs for each condition are shown in the set size rows on the left of the plot. The vertical bars represent the number of DEGs unique to the subset indicated by the colored dots in the intersection matrix. Intersections showing genes that are common to SD regardless of genotype and common to RS regardless of genotype are shown.
