## Additional File 8 for "Shank3 mutation disrupts the molecular signature of sleepiness across development"

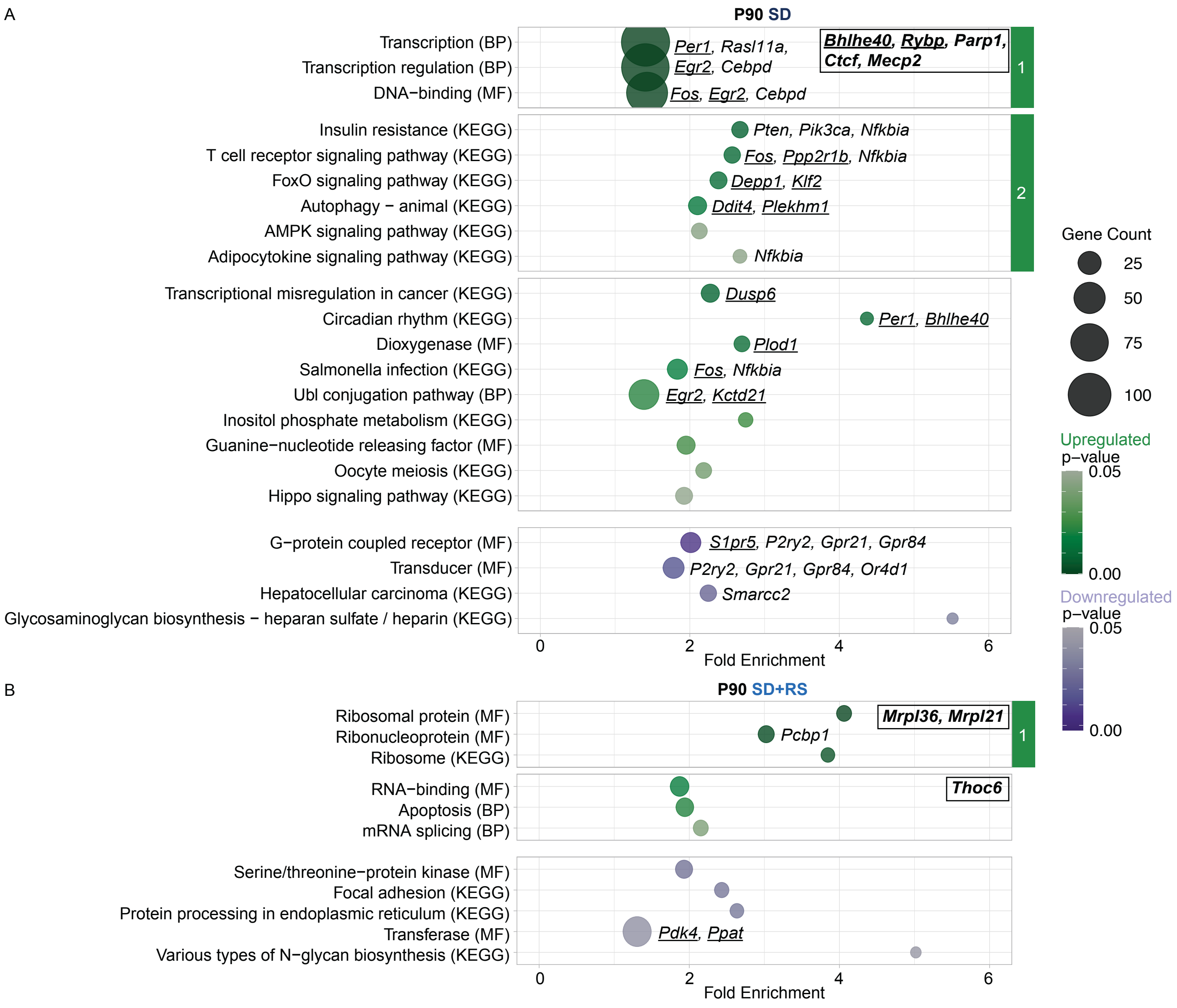


**Additional File 8**. **Sleep manipulations impact DNA damage detection, circadian rhythms, stress responses regardless of genotype at P90.** Functional enrichment analysis plots for DEGs common to **A**) 5 hours of SD and **B**) 5 hours of SD followed by a 2-hour of recovery period. Enriched functional annotation terms (modified Fisher's exact *P* value < 0.05) from UniProt BP, MF and KEGG are displayed vertically. Circle size indicates the number of genes found within a given term which are plotted based on fold enrichment as displayed on the x-axis. Upregulated terms are in green and downregulated terms are in purple with darker shades representing smaller *P* values. Hub genes are bolded and displayed in boxes in the upper right corner of their corresponding cluster. . Enrichment scores for each cluster in **A**: upregulated cluster 1 (4.42), upregulated cluster 2 (1.76). Enrichment scores for each cluster in **B**: upregulated cluster 1 (2.74).
