## Additional File 10 for "Shank3 mutation disrupts the molecular signature of sleepiness across development"

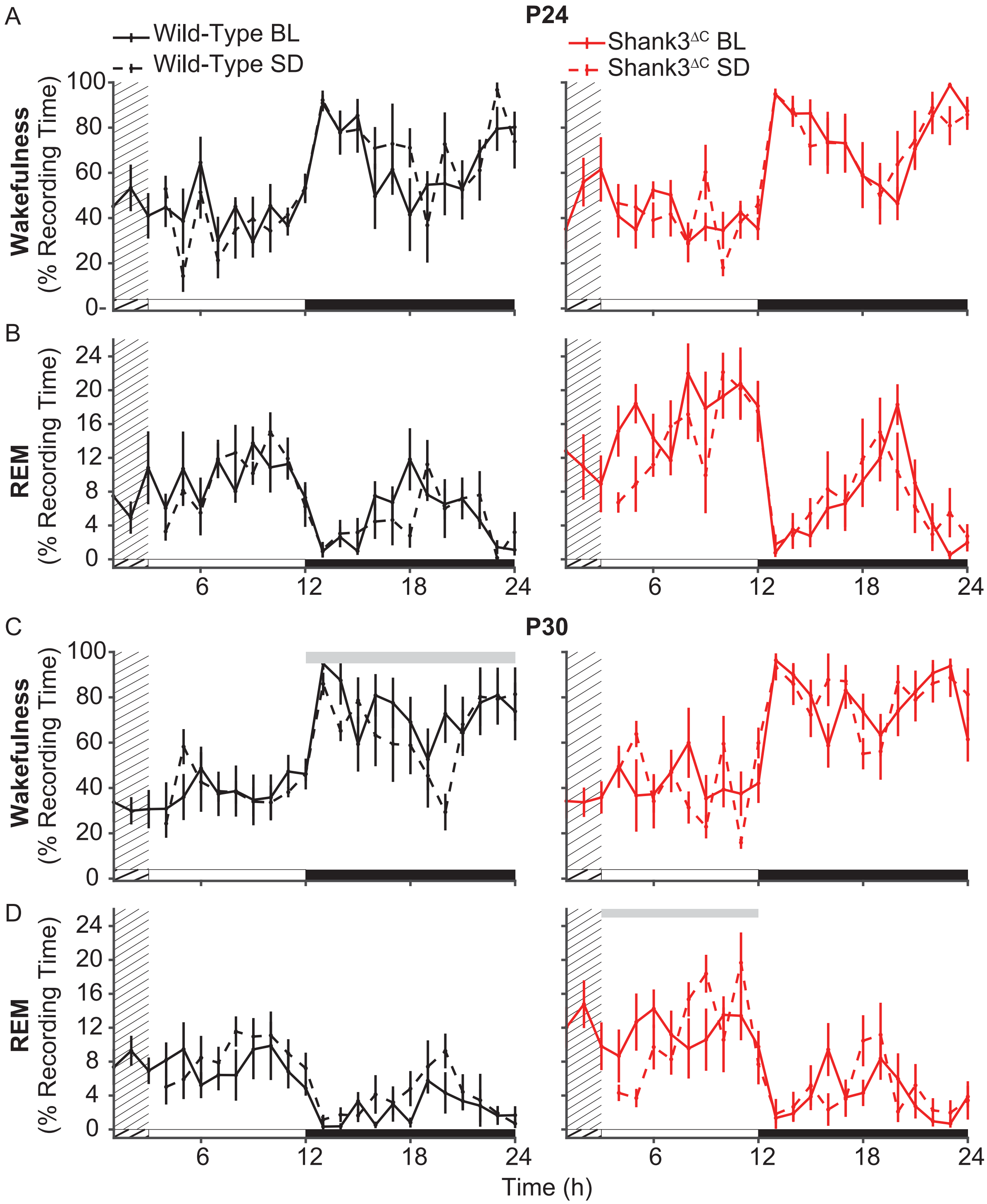


**Additional File 10. WT animals are awake less following SD at P30 while Shank3$^{\Delta C}$ animals show differences in REM sleep.** Total recording time (TRT) in wake (top row) and REM sleep (bottom row) during recovery sleep compared to baseline. **A-B**) P24. **C-D**) P30. The gray bar above the plots represents significance from repeated measures ANOVA across hour 3-12 (light period) and 13-24 (dark period) ($p < 0.05$). Light period is represented by the white bar, dark period by the black bar. The solid line represents the baseline day and the dashed line represents the recovery sleep day with the grated area representing 3 hours of SD. WT animals are represented in black (n = 5 P24, 6 P30) and Shank3$^{\Delta C}$ animals are represented in red (n = 6 P24, 6 P30).
