## Additional File 17 for "Shank3 mutation disrupts the molecular signature of sleepiness across development"

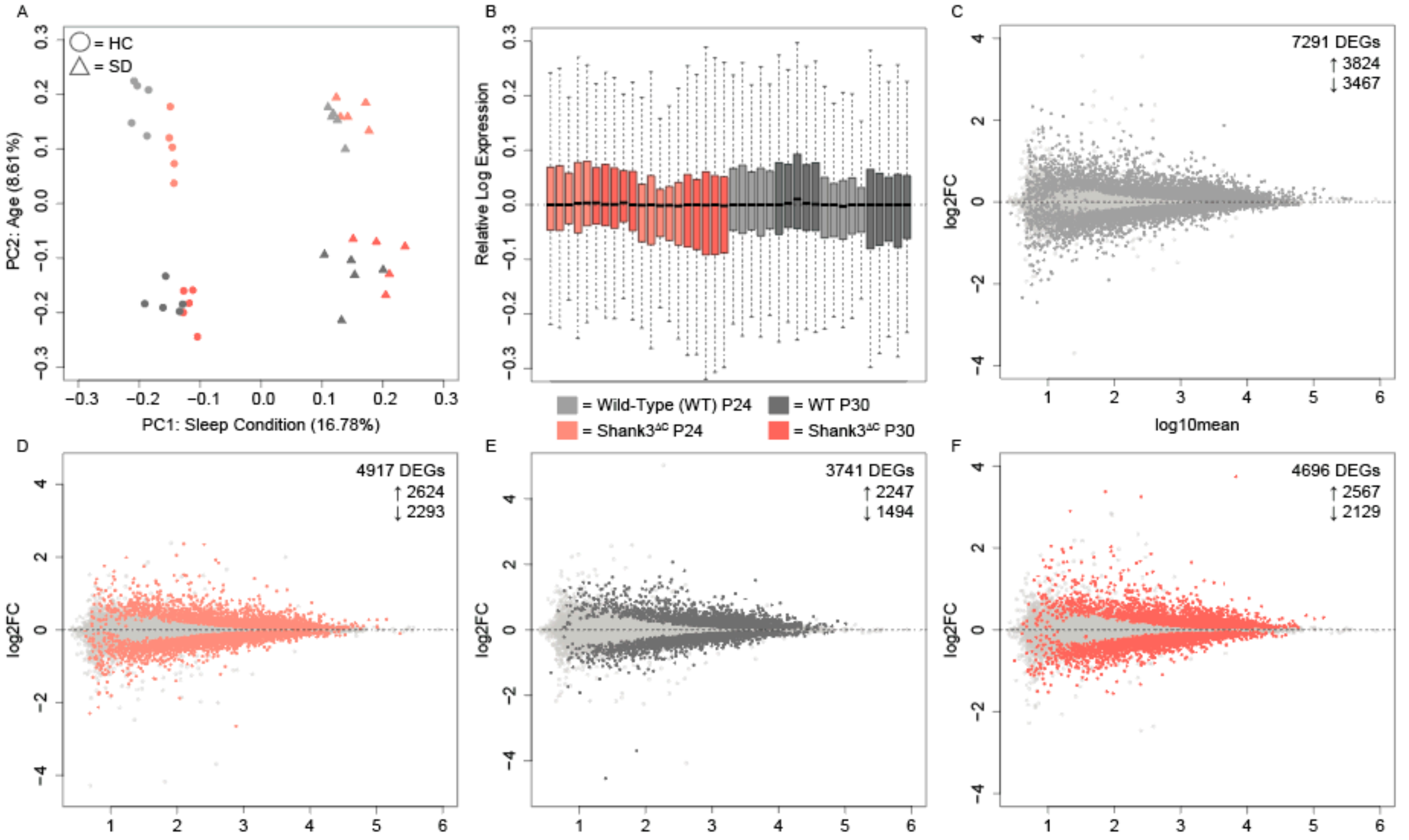


**Additional File 17. WT and Shank3$^{\Delta C}$ gene expression differs following SD at P24 and P30. A**) Principal component analysis following RUVs normalization with k = 15 unwanted factors. Circles represent home cage (HC) animals and triangles represent SD animals. WT animals are shown in shades of grey and Shank3$^{\Delta C}$ animals are shown in shades of red. n = 5 for each condition. Sleep condition (PC1, 16.78%) and age (PC2, 8.61%) account for the most variability within the data. **B**) Relative log expression for each sample and condition. Color code as in A. **C-F**) MA plots following differential expression analysis on RUV normalized gene counts. **C**) WT postnatal day 24 (P24), **D**) S3P24, **E**) WT postnatal day 30 (P30), **F**) S3P30. Color code as in A showing differentially expressed genes (DEGs) in light grey for P24 WT animals, medium grey for P30 WT animals, light red for P24 Shank3$^{\Delta C}$ animals and medium red for P30 Shank3$^{\Delta C}$ animals. Numbers of total DEGs, upregulated DEGs, downregulated DEGs are reported in the top right corner for each condition.
